# Mitotic BLM functions are required to maintain genomic stability

**DOI:** 10.1101/2025.06.16.659902

**Authors:** Tamara Eleanore Hamann, Angela Wieland, Andrea Tirincsi, Kruno Vukušić, Farbod Mohseni, Rene Wardenaar, Marialucrezia Losito, Ipek Ilgin Gönenc, Bernd Wollnik, Floris Foijer, Iva M. Tolić, Zuzana Storchova, Markus Räschle

## Abstract

The BLM helicase is a critical genome maintenance protein involved in diverse cellular processes including DNA replication, repair, transcription, and chromosome segregation. During mitosis, it cooperates with the PICH helicase and topoisomerases to resolve ultrafine DNA bridges (UFBs) - non-chromatinized DNA structures that link sister chromatids - through a mechanism that is not yet fully understood. Here we tagged endogenous BLM and PICH with fluorescent proteins and BLM with an auxin-inducible degron to generate a cell model system that enables temporal tracking of UFB dynamics in the presence or absence of BLM. Time-resolved lattice light sheet microscopy established the dynamic localization patterns of BLM and PICH throughout the cell cycle. While BLM cycles between PML bodies and DNA repair foci in interphase, it dissociates from chromatin at the mitotic entry, and re-associates during anaphase to UFBs as well as to CENP-B-positive mitotic foci. Acute BLM depletion during mitosis increased the fraction of unresolved UFBs, micronuclei containing acentric fragments, binucleation, and resulted in subtle genomic abnormalities detected by single-cell whole genome sequencing. These findings highlight a mitosis-specific role for BLM in UFB resolution and underscore its function in preserving genomic stability.

**Graphical abstract:** 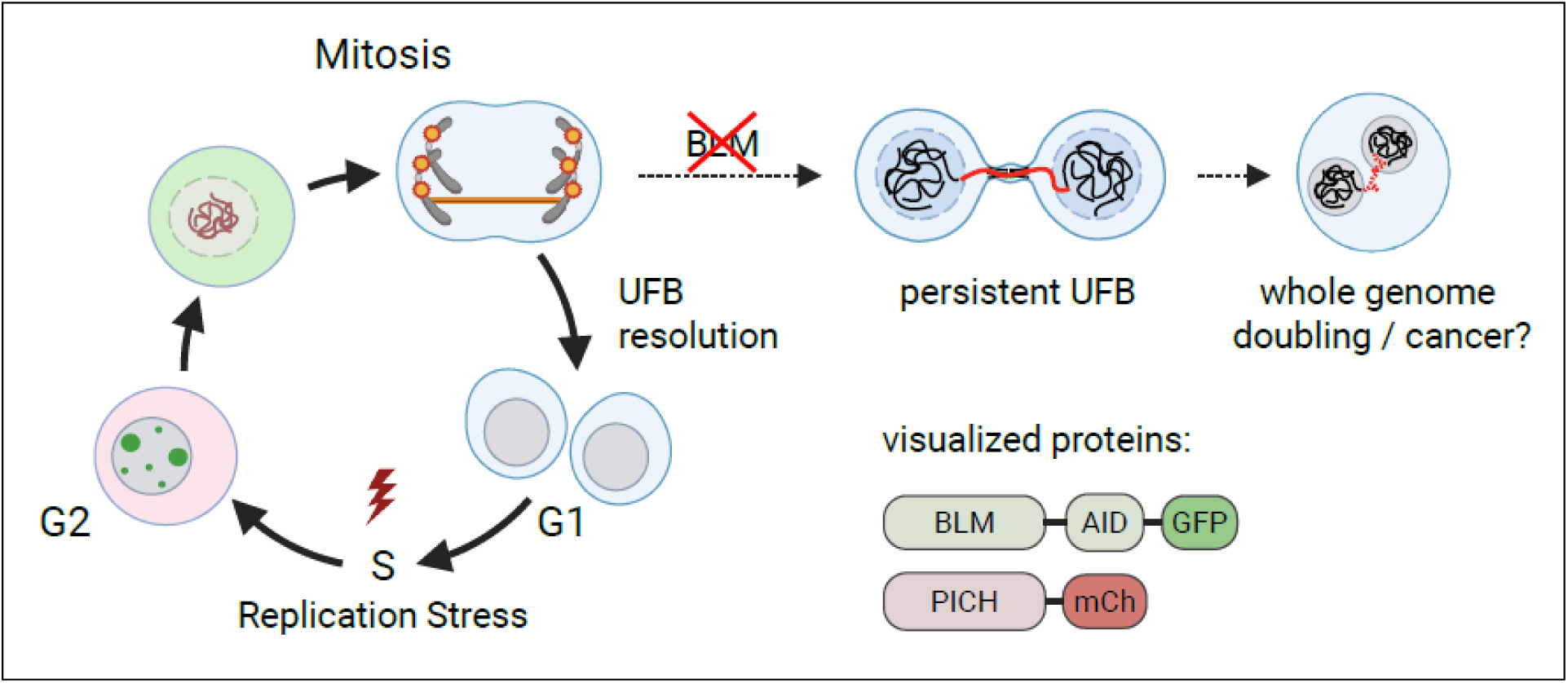

## Introduction

Bloom syndrome is a rare genetic disorder characterized by developmental defects, immunodeficiency, and a strikingly high incidence of early-onset cancers (1). It is caused by bi-allelic mutations of the *BLM* gene (2). Cells of Bloom syndrome patients show features of increased genomic instability, such as elevated rates of sister chromatid exchanges, mitotic aberrations, and chromosome breakage. The *BLM* gene encodes a highly conserved 3’ – 5’ ATP-dependent helicase and its expression is tightly regulated throughout the cell cycle. The abundance sharply increases at the beginning of S phase, reaching maximum levels during S and G2 phase. In late mitosis, BLM is degraded by the ubiquitin-proteasome system and remains undetectable during most of the G1 phase (3,4). BLM protein associates with topoisomerase 3 alpha (TOP3A) and RecQ-mediated genome instability 1 and 2 (RMI1 and RMI2) in the so-called BTRR complex; inactivation of these interactors causes the Bloom syndrome-like disorder (5,6). Together, they promote strand resection as well as dissolution of recombination intermediates. In addition, BLM has been reported to facilitate the restart of stalled replication forks and the removal of secondary structures, such as G-quadruplex, in DNA and RNA (reviewed in (7,8)).

Consistent with its multiple functions in various DNA metabolic pathways, the subcellular localization of the BLM helicase and its interaction with other proteins are highly dynamic and tightly regulated. BLM primarily localizes to membrane-free, nuclear structures called promyelocytic leukemia nuclear bodies (PML-NB) (9,10), as well as to diffuse patches representing the nucleolus, where it likely contributes to the replication and transcription of the repetitive rDNA sequences (11,12). In S and G2 phase cells, DNA damage triggers the rapid relocalization of BLM from PML-NBs into small DNA repair foci positive for phosphorylated H2AX, and several other DNA repair proteins including RAD51 and FANCD2(13,14). A subfraction of these repair foci also stains positive for the single-stranded DNA-binding replication protein A (RPA), and proliferating-cell-nuclear-antigen (PCNA), suggesting they form at sites of stalled DNA replication forks (15–17). As cells enter mitosis, BLM becomes highly phosphorylated by PLK1, MPS1, CDK1 and other mitotic kinases, which regulates its interaction with other proteins and DNA (4,18,19). In anaphase, BLM co-localizes with polo-like kinase 1 (PLK1)-interacting helicase (PICH) at so-called ultrafine bridges (UFBs) (20) that physically link the segregating sister chromatids. Transient UFBs are sporadically observed in untreated, healthy cells, indicating that their resolution is a normal physiological process essential for accurate sister chromatid segregation. Interference with DNA replication, or the induction of homologous recombination strongly increase UFB formation (21). Accordingly, UFBs are often formed within difficult-to-replicate genomic regions, or in hotspots of mitotic DNA recombination, such as common fragile sites, rDNA, telomeres, or centromeres (22,23). Although the exact structure of the DNA residing in UFBs remains largely unknown, different classes of UFBs have been proposed to be composed of either late replication intermediates consisting of double stranded DNA flanked by stalled replication forks, fully replicated DNA held together by catenanes or recombination intermediates, respectively (recently reviewed in (24)). UFBs are free of nucleosomes and cannot be visualized by standard DNA dyes, such as DAPI or Hoechst, suggesting that the DNA between the segregating sisters is either highly stretched or single-stranded (25). Instead, UFBs can be identified by associated proteins, including the PICH and BLM helicases. The current model for the resolution of replication-stress induced UFBs involves the recognition of stretched DNA by the PICH helicase, followed by the recruitment of the BTRR complex and other proteins to the UFB. Subsequently, the BLM helicase unwinds double stranded DNA, while TOP3A and TOP2A help to resolve topological structures to unlink the sister chromatids (26–29). Taken together, the BLM helicase not only plays a crucial role in genome maintenance during S and G2, where it promotes DNA replication and recombination, but also in mitosis, where it supports effective UFB disjunction. However, to what extent the early and late functions of BLM contribute to genome stability remains largely unknown. Here we established a cell line allowing *in vivo* visualization of UFBs through fluorescently tagged BLM and PICH. Using this cell line, we demonstrate that BLM dissociates from chromatin at nuclear envelope breakdown and is subsequently recruited *“de novo”* to UFBs during anaphase. Moreover, BLM accumulates at anaphase onset in previously uncharacterized “mitotic foci”, frequently localized near centromeres. BLM depletion stabilizes UFBs and promotes genomic instability. Notably, mitosis-specific BLM depletion underscores its essential role in UFB resolution and genome maintenance during cell division.

## Material and Methods

### Antibodies

| Antibody | Company | Dilution | Application<sup>1)</sup> |
| --- | --- | --- | --- |
| Mouse anti-53BP1 | Millipore, MAB3802 | 1:500 | IF |
| Mouse anti-α actinin | Santa Cruz, sc-17829 | 1:1000 | WB |
| Rabbit anti-γH2Ax | Abcam, ab2893 | 1:500 | IF |
| Mouse anti-BLM | Santa Cruz, sc-365753 | 1:250 | WB |
| Mouse anti-CENP-B | Santa Cruz, sc-376392 | 1:1000 | IF |
| Rabbit anti-Cyclin A2 | Abcam, 181591 | 1:500 | IF, WB |
| Mouse anti-Cyclin B1 | Cell Signaling, 4135 | 1:2000 | WB |
| Rat anti-GFP | ChromoTek, 3H9 | 1:250 | WB |
| Rabbit anti-mCherry | Proteintech, 26765-1-AP | 1:1000 | WB |
| Mouse anti-MPM2 | Merck, 05-368 | 0.06 μg | FC |
| Rabbit anti-PICH | Abnova, H00054821-D01 | 1:1000 | WB |
| Rabbit anti-PICH | Abnova, H00054821-D01 | 1:150 | IF |
| Mouse anti-PML | Santa Cruz, sc-966 | 1:250 | IF |
| Rabbit-anti-RAD51 | Santa Cruz, sc-8349 | 1:250 | IF |
| Alexa Fluor 594 donkey, anti- mouse | Biozol, JIM-715-585-150 | 1:500 | IF |
| Alexa Fluor 647 donkey, anti-rabbit | Biozol, JIM-711-605-152 | 1:500 | IF |
| Alexa 488-conjugated goat anti-mouse IgG | Biozol, JIM- 115-545-003 | 1:300 | FC |
| Anti-mouse HRP | R&D Systems, HAF007 | 1:5000 | WB |
| Anti-rabbit HRP | R&D Systems, HAF008 | 1:5000 | WB |
| Anti-rat HRP | R&D Systems, HAF005 | 1:5000 | WB |

<sup>1)</sup> IF: Immunofluorescence; WB: Western Blotting; FC: Flow Cytometry

### Cas9/sgRNA plasmid construction

The Cas9/sgRNA plasmid was constructed using the px330-U6-Chimeric_BB-CBh_hSPCas9 plasmid obtained from Feng Zhang (Addgene plasmid #42230)(30). Two oligonucleotides, each containing a target-specific guide RNA sequence of 20 nucleotides and appropriate overhangs, were cloned into the px330-based plasmid. The target-specific guide RNA sequence was selected using the Benchling tool, prioritizing sequences proximal to the stop codon with optimal off-target and on-target scores for c-terminal gene tagging.

### Homology donor plasmid construction

Homology donor plasmids were generated through a multi-step cloning process. First, target-specific homology arms, fluorescence markers, the 44–amino acid version of the AID tag (31) and the resistance genes were individually cloned into the linearized pIDC backbone plasmid (Geneva Biotech) to generate the pENTRY constructs using the Gibson Assembly Cloning Kit (NEB #E2611). Subsequently, homology donors were generated by combining the inserts of the pENTRY vectors with the acceptor vector pSG3651 using the NEBridge Golden Gate Assembly Kit (NEB #E1602), and error-free assembly was confirmed by DNA sequencing. The DNA insert sequences for the individual homology arms (800 bp), the AID domain, the fluorescence marker enhanced green fluorescence protein and mCHERRY were ordered as gblocks with corresponding overhangs for Gibson assembly from Integrated DNA Technologies (see Fig. 1 and Supplementary Fig. 1 and 5 for details). Inserts for the hygromycin and neomycin resistance genes were PCR amplified from pMK292 (Addgene #72830) and pMK293 (Addgene #72831). All sequences are available upon request.

**Figure 1.**
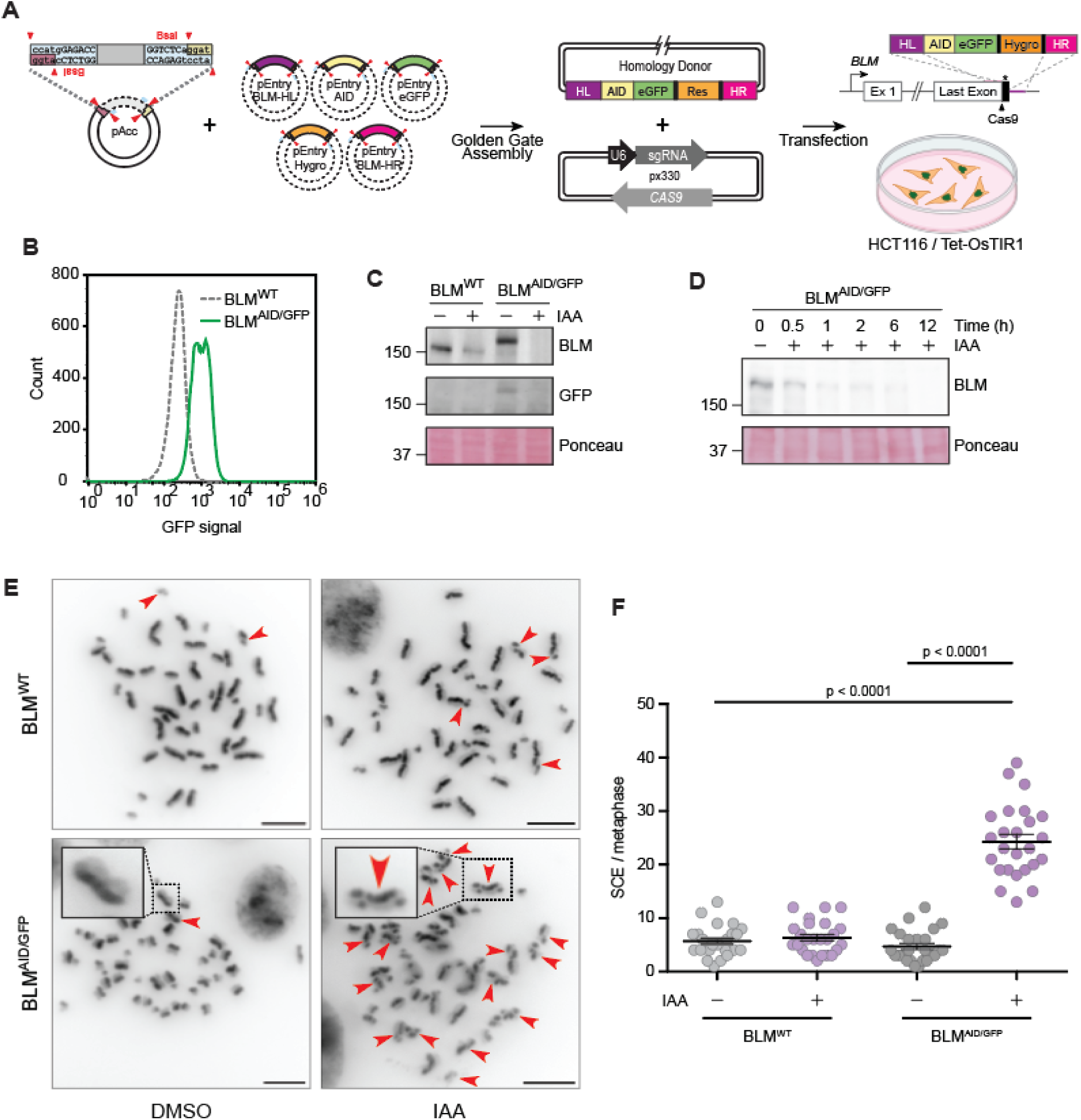
Endogenous AID-eGFP tagging of BLM facilitates its rapid degradation. **(A)** Schematic illustration of the gene editing strategy. (**B**) Flow cytometry showing the GFP signal in parental HCT116 and tagged BLM^AID/GFP^ cells. (**C-D**) Immunoblotting of the lysates from parental HCT116 and BLM^AID/GFP^ cells with and without auxin treatment (indole-3-acetic acid, IAA) for 24 h (C) or as indicated (D). Proteins were detected with the indicated antibodies, Ponceau S staining served as loading control. (**E**) Sister chromatid exchange (SCE) visualized on metaphase spreads. Asynchronous HCT116 or BLM^AID/GFP^ cells were incubated with IAA or DMSO for two rounds of replication. Red arrows point to metaphase chromosomes containing SCEs; insets show representative examples. (**F**) Quantification of SCE. 25 metaphase spreads were quantified for each condition, each dot represents an individual metaphase spread; mean with S.E.M., Student t-test.

### Cell culture, transfection, and colony isolation

The parental human colon cancer cells, HCT116 Tet-OsTIR1 ((32), a kind gift from Dr. Masato Kanemaki), were cultured in Dulbecco’s Modified Eagle Medium (DMEM) with high glucose and GlutaMAX (Thermo Fisher Scientific). The medium was supplemented with 10 % fetal bovine serum (FBS) (Thermo Fisher Scientific), 1 % streptomycin/penicillin (Thermo Fisher Scientific), and 1 µg/mL puromycin (Thermo Fisher Scientific). The cells were maintained in a 5 % CO2 humidified incubator at 37 °C. Cells were routinely tested for the absence of mycoplasma contamination (MycoStrip, Invivogen). Plasmids and homology donors were co-transfected with PEI (Polyscience) for 6 h and after two days, the cell suspension was distributed onto 15 cm culture plates to ensure single-cell distribution, followed by selection in medium containing 100 µg/mL hygromycin B or neomycin (both Invivogen). After 14 days, individual colonies were picked, expanded, and validated by PCR, flow cytometry and Western blotting. For PICH tagging, neomycin resistant, mCherry-positive single cells were sorted into individual wells of a 96 well plate using a SONY SH800S cell sorter and expanded in the presence of neomycin. Successful PICH tagging was validated by Western Blotting and Sanger sequencing. In all experiments, cells were harvested by trypsinization. Cells were centrifuged at 1300 rpm for 3 min, cell pellets were washed once with PBS and directly processed, as indicated.

### Auxin-induced protein degradation and cell cycle synchronization

To induce degradation of the AID-tagged BLM protein, HCT116-Tet-OsTIR1 BLM^AID/GFP^ were preincubated for at least 24 h with 1 µg/ml doxycycline to induce expression of OsTIR1, followed by incubation with 500 μM auxin (indole-3-acetic acid, IAA). For cell synchronization, cells were arrested at the G1/S boundary by addition of thymidine (2 mM final, 24 h). To release cells from the single thymidine block, cells were washed trice with warm PBS and placed into fresh media containing 100 nM APH to induce mild replication stress or DMSO as a control. For continuous BLM depletion (“S+M” depletion), auxin was added together with thymidine at the start of the cell synchronisation. To restrict BLM depletion to mitosis (“M”), auxin was added at the indicated time after the release from the thymidine block. After the thymidine wash-out, media was replenished with fresh doxycycline (1 ug/ml) and auxin (500 μM) as indicated.

For scWGS analysis, 1 x 10^6^ BLM^AID/GFP^ cells were seeded per 10 cm dish and synchronized as described above, except that the cells were released into fresh media containing 10 µM EdU to monitor progression through S phase. 6 h after the release from the thymidine block, cells were washed thrice with PBS and incubated in media containing 100 nM APH to induce mild replication stress and 500 nM palbociclib (Seleck) to arrest the cells at the next G1 phase. After 18 h cells were harvested and fixed using 70 % ethanol for flow cytometry (see below). In a parallel experiment EdU was omitted and cells were sorted into single wells for scWGS analysis (see below).

### Flow cytometry

To validate the expression of GFP-tagged BLM, asynchronous cells treated for the indicated time with IAA or ETOH were harvested. Cell pellets were resuspended in 200 μL of PBS and analysed using the Attune™ NxT Flow Cytometer (Thermo Fisher Scientific) equipped with a 488 nm blue excitation laser and a 530/30 bandpass filter for GFP emission detection. Data acquisition and analysis were performed using FlowJo software v. 10.1 (Becton, Dickinson and Company). During analysis, cell debris and cell doublets were excluded based on forward versus side scatter parameters.

To analyse cell cycle progression, pellets of synchronized cells harvested at the indicated time points were fixed by slowly adding 500 µl of 70 % ice-cold ethanol while vortexing, followed by overnight incubation at 4 °C. The next day, ethanol was removed by washing the cells with PBS containing 0.25 % Triton-X-100, and centrifuged at 2000 rpm for 5 min. For immunostaining, cells were incubated for 1 h at RT with 0.06 μg MPM-2 monoclonal antibody resuspended in 100 µl PBS containing 1 % BSA. After washing, cells were incubated for 30 min at RT in the dark with Alexa 488-conjugated goat anti-mouse IgG antibody diluted 1:300 in 100 μl PBS containing 1 % BSA. Subsequently, cells were stained with 500 μl of FxCycle™ PI/RNase staining solution (Themo Fisher Scientific) and incubated for 30 min at RT in the dark. Measurements were taken using the Attune™ NxT Flow Cytometer (Thermo Fisher Scientific) with 488 nm blue excitation and a 530/30 bandpass filter for emission detection of the MPM2 signal and 574/26 bandpass filter for the DNA stain propidium iodide (PI) emission detection. Data was analyzed as described above and the fraction of cells in G1, S, G2 and M phase was quantified by applying gates to the PI and histogram plot.

To measure EdU incorporation, cells were permeabilized for 15 min with Fix/Perm solution (Thermo Fisher Scientific) according to the manufacturer’s instructions. After permeabilization, 1 ml PBS was added, and the samples were centrifuged for 3 min at 1300 rpm. The supernatant was discarded, and the cell pellets were resuspended in 100 µl Perm wash (Thermo Fisher Scientific). Subsequently, 500 µl Click-iT Reaction Mix (1µM Eterneon Red (Baseclick GmbH), 6.6 % (v/v) 1.5 M Tris (pH 8.8), 500 µM CuSO4, 100 mM Ascorbic Acid in PBS) was added to the samples for 20 min at room temperature in the dark. After the incubation, the samples were washed three times with 500 µl Perm wash (centrifugation for 2 minutes at 1600 rpm). Finally, for DNA staining cells were incubated in 300 – 500 µl PBS containing 1 µg/ml DAPI and 10 µg/ml RNase. Flow cytometry was performed with an Attune™ NxT Flow Cytometer. DAPI and EdU-bound Eterneon-Red were excited by a 405 nm and a 638 nm laser, respectively. The emission from the excited DAPI was collected with a 440/50 filter. The light emitted by the excited Eterneon-Red was collected with a 670/14 filter. Data was analysed as described above and EdU incorporation was quantified by plotting the Eterneon-Red against the DAPI intensity.

### Sister Chromatid Exchange (SCE) Assay

To label sister chromatids, cells were incubated for two rounds of cell division (40 h) with 20 µM bromodeoxyuridine. At a confluency of 70–80 %, cells were treated with 400 ng/ml colchicine for 5 h prior harvesting. Cells were collected by trypsinization and centrifuged at 1200 rpm for 3 min. Cell pellets were resuspended in 75 mM KCl and incubated for 15 min at 37 °C. Fixation was performed with 3:1 methanol/acetic acid for three rounds. Fixed samples were dropped onto glass slides. After drying of the spreads overnight at RT, slides were immersed in 10 µg/ml Hoechst 33258 for 20 min. After rinsing the slides in 1X Sorensen phosphate buffer (50 % 0.1 M Na_2_HPO_4_, 50 % 0.1 M KH_2_PO_4_, pH 6.8), slides were exposed to UV-A for 30 min. Slides were incubated in 1x salt sodium citrate buffer (3 M NaCl, 300 mM sodium citrate, pH 7.0) for 1 h at 50 °C. Slides were further immersed in 4 % Giemsa stain for 30 min at RT. Vectashield Mounting Medium with DAPI was used to visualize the DNA. Slides were washed subsequently in H_2_O twice and dried overnight at RT in the dark. 25 metaphases were captured per condition. Microscopy was performed using the 63x oil objective of a semi-automated inverted Zeiss microscope (AxioObserver Z1). Images were acquired using SlideBook 6 (Intelligent Imaging Innovations).

### Quantification of DNA breaks on mitotic chromosome spreads

Synchronized cells were collected 10.5 h after the release from a single thymidine block (see above). Cell pellets were resuspended in 75 mM KCl and incubated for 15 minutes at 37 °C. Fixation was performed by resuspending the cell pellets with 3:1 methanol/acetic acid for three rounds. Fixed samples were dropped onto glass slides. Vectashield Mounting Medium with DAPI was used to visualize the DNA. 25 metaphases per replicate from 3 independent experiments were captured. Microscopy was performed using the 63x oil objective of a semi-automated inverted Zeiss microscope (AxioObserver Z1). Images were acquired using SlideBook 6 (Intelligent Imaging Innovations).

### Protein isolation, SDS-PAGE and Western blotting

For whole cell lysates, cell pellets were resuspended in RIPA buffer (50 mM Tris-HCl, pH 7.5, 150 mM NaCl, 1 % NP-40, 0.5 % sodium deoxycholate, 0.1 % SDS, 1 mM EDTA) supplemented with protease inhibitors (complete EDTA-free, Roche) and phosphatase inhibitors (Roche) and sonicated. The lysate was incubated for 25 minutes at 4 °C. Protein concentration was determined using the Pierce BCA Protein Assay Kit (ThermoFisher, #23225). Equal amounts of protein were resuspended in 1x Laemmli buffer (62.5 mM Tris-HCl (pH 6.8), 2 % SDS, 10 % glycerol, 0.01 % bromphenol blue, 1 % β-mercaptoethanol), denatured for 5 min at 95 °C and separated by sodium dodecyl sulfate-polyacrylamid gel electrophoresis. Proteins were transferred onto a nitrocellulose membrane (GE Healthcare). Protein transfer was validated by Ponceau S staining. Next, membranes were blocked for 1 h in 3 % BSA in TBS (only for detecting BLM) or with 5 % skim milk in TBST (Tris-buffered saline with 0.1 % Tween 20) TBS-T, followed by overnight incubation at 4 °C in primary antibodies. On the next day, the membranes were washed and incubated for 1 h in secondary antibodies at RT. Proteins were visualized with enhanced chemiluminescence (ECL) reagent (BioRad) and scanned on an Azure c300 imaging system (Azure Biosystems). Protein normalization was performed using housekeeping proteins. Signal intensities were quantified using the software Fiji (National Institutes of Health).

### Immunofluorescence staining

Cells were cultured on cover slips placed in 6-well plates. The cells were fixed with ice-cold 100 % methanol for 10 min on ice or with 4 % PFA (for mitotic cell analysis) for 20 minutes at room temperature. Both fixing reagents were removed by washing the cells thrice with PBS. To permeabilize the cells, 0.5 % Triton 100-X (diluted in PBS) (Carl Roth) was added for 20 minutes. Afterwards, the cells were washed thrice with PBS, and then blocked (3 % BSA, 1 % NaN3, diluted in PBS-T) for 1 h at room temperature. To stain for the protein of interest, the cells were incubated with the respective primary antibodies overnight at 4 °C. The following day, the cells were washed thrice with PBS and then incubated in the appropriate secondary antibody, for 1 h in the dark at room temperature. After removing unbound secondary antibodies by washing with PBS, the DNA was stained by incubating the cells for 5 minutes with 0.2 μg/ ml DAPI (Carl Roth), diluted in PBS. For BLM visualization, the endogenous GFP signal was detected at a wavelength of 488 nm without antibody staining. The cells were washed once with PBS and once with water before mounting the coverslips on glass slides. Images were captured using the 40x or 60x oil objective of the Eclipse Ti2-E with Spectra III epifluorescence microscope (Nikon). For the detection of anaphase cells, z-stacks were captured with relative offsets with 25 steps and a size of 0.3 µm, range 7.2 µm. The images were analysed manually using the software Fiji.

### Live cell imaging

Live-cell imaging was performed as described in (33). Multiple 3.5 cm glass-bottom dishes were seeded at various densities. Cells that reached approximately 70 % confluency were chosen to be imaged. 12 1 s before imaging start the cells were stained with 1 nM of the nuclear stain SPY555-DNA (λ_abs_ 555 nm) at 37 °C. Additionally, 1 μM Verapamil was added to prevent dye efflux. The cells were placed in a humidified chamber with 5 % CO_2_ at 37 °C. Images were taken every 2 minutes for a minimum of 12 h using Lattice Light Sheet 7. The images were analysed manually using the software Fiji. For tracking individual cells or foci the Fiji plugin TrackMate was used.

### Statistical analysis

Data were analyzed using GraphPad Prism (version 9.0). All fluorescent images were blind counted and statistical comparisons between two groups were performed using an unpaired two-tailed Student’s t-test. P-values of less than 0.05 were considered statistically significant and are shown on the graphs. The number of independent biological replicates (n) and the total number of cells or events analyzed are indicated in the figure legends. For protein co-localization analysis, Pearson correlation coefficients were calculated using the Fiji plugin Coloc 2.

### scWGS and AneuFinder analysis

Sequencing was performed using a NextSeq 500 machine (Illumina; up to 77 cycles single end). The generated data were on beforehand demultiplexed using sample-specific barcodes and changed into fastq files using bcl2fastq (Illumina; version 1.8.4). Reads were afterwards aligned to the human reference genome (GRCh38/hg38) using Bowtie2 (version 2.2.4; (34)). Duplicate reads were marked with BamUtil (version 1.0.3;(35)). The aligned read data (bam files) were analysed with a copy number calling algorithm called AneuFinder (version 1.14 and 1.30; https://github.com/ataudt/aneufinder; (36)). Following GC correction and blacklisting of artefact-prone regions (extreme low or high coverage in control samples), libraries were analysed using the dnacopy and edivisive copy number calling algorithms with variable width bins (average binsize = 1 Mb; step size = 500 kb). Blacklists and variable width bins were constructed with the use of an euploid reference (37). Results were afterwards curated by requiring a minimum concordance of 90 % between the results of the two algorithms. Libraries with on average less than 10 reads per chromosome copy of each bin (2-somy: 20 reads, 3-somy: 30 reads, etc.) were discarded. This minimum number of reads comes down to roughly 55,000 for a diploid genome. The copy numbers of the T24 samples (T24 CTR, T24 M, T24 S+M) were expressed relative to the median (consensus) copy number profile of the T0 control sample (T0 CTR) in order to account for the non-euploid nature of this cell line. This median copy number was determined for each bin (median across libraries). The copy numbers of this consensus copy number profile were doubled when compared to tetraploid libraries.

### Calculation scores

Aneuploid scores for each sample were first calculated for each library by calculating the average absolute difference from the median copy number profile of the T0 control sample. The size of the bins was used as weight (weighted average). The genome-wide scores were calculated as the average of all the libraries of the sample. The heterogeneity score of each bin was calculated as the proportion of pairwise comparisons (cell 1 vs. cell 2, cell 1 vs cell 3, etc.) that showed a difference in copy number (e.g. cell 1: 2-somy and cell 2: 3-somy). The heterogeneity score of each sample was calculated as the weighted average of all the bin scores (size of the bin as weight). The structural score of each library was simply calculated as the average number of copy number transitions (break points) divided by one million. The genome-wide scores were calculated as the average of all the libraries of the sample. For fragment size analysis copy number segments with a maximum size of 10 Mb were selected. The segments were required to show a copy number that is different from the median (consensus) copy number.

## Results

### Rapid degradation of BLM impairs genome stability

To study the distinct functions of BLM during different cell cycle phases, we modified the endogenous *BLM* by a c-terminal fusion with an auxin inducible degron (AID) domain for rapid degradation of the protein (38). In addition, eGFP was included for tracking the endogenous BLM protein in live cells. CRISPR/Cas9 knock-ins require homology donors, in which inserts are flanked by two homologous regions matching the genomic sequence at the target site. In HCT116 cells, homology arms of at least 500 base pairs (bp) are required for efficient tagging (39). To this end, we developed a modular cloning system, in which targeting vectors can be assembled from interchangeable parts by PCR-free Golden Gate Assembly (Fig. 1A, Supplementary Fig. 1A and B). This allows to easily combine a degron domain with different fluorescence markers (e.g., eGFP, mCherry) and resistance genes (e.g., hygromycin (Hygro^R^) or neomycin (Neo^R^)). For BLM tagging, a homology donor was assembled from the AID, eGFP and Hygro^R^ entry vectors joined by two flanking homology arms of 800 bp matching the sequence to the left and right of the stop codon in the endogenous *BLM* gene (Fig. 1A). CRISPR/Cas9 mediated gene editing was carried out with a single guide RNA targeting the Cas9 endonuclease to cut 12 nucleotides upstream of the *BLM* stop codon (Supplementary Fig. 1C). A plasmid expressing the guide RNA and Cas9 was transiently transfected into HCT116-OsTIR1 cells together with the homology donor (Fig. 1A, Supplementary Fig. 1D). Flow cytometry and Western blotting with antibodies against GFP and BLM confirmed successful homozygous tagging in one clone, here referred to as BLM^AID/GFP^ (Fig. 1B and C). Exposure of asynchronous cells to auxin for 24 h reduced the levels of the AID-tagged BLM protein to below detectable levels (Fig. 1C) and time-course experiments revealed a rapid depletion of BLM with its levels dropping to approximately 10 % within 1 h after auxin addition (Fig. 1D). Loss of BLM is associated with a hallmark genomic instability phenotype known as sister chromatid exchange (SCE)(40). Consistently, the addition of auxin to the BLM^AID/GFP^ cells increased the rate of SCE events 5-fold compared to untreated control cells (Fig. 1E and F). Importantly, in the absence of auxin, SCE levels were comparable to those observed in the parental HCT116 cells with or without auxin treatment. We conclude that AID-mediated depletion of BLM mimics the phenotype of BLM loss observed in cells from patients with Bloom syndrome. Furthermore, the results confirm that the AID/GFP-tagged BLM protein is fully functional and efficiently suppresses the resolution of recombination intermediates that give rise to sister chromatid exchanges.

### Dynamic localization of the BLM helicase in mitotic cells

First, we examined BLM localization dynamics during the cell cycle. To this end, we performed live cell imaging of asynchronous BLM^AID/GFP^ cells with captures every 2 minutes using Lattice light sheet microscopy (33). To induce mild replication stress, cells were incubated with the polymerase inhibitor aphidicolin (APH) at a low concentration of 100 nM for 12 h before imaging the live cells to ensure that all captured mitotic BLM^AID/GFP^ cells had passed through a perturbed S phase (Fig. 2A). Notably, the low dose of APH induced mild replication stress without triggering a cell cycle arrest (41). Live cell imaging revealed a recurrent pattern of the BLM-GFP protein on chromatin (Fig. 2B, Movie M1A and B). In the G2 phase, before the chromosome condensation started, we observed a diffused chromatin signal of BLM-GFP, as well as several more prominent foci (Fig. 2B). The foci were confirmed to largely represent PML-positive nuclear bodies (PML-NB), as the BLM signal strongly co-localized with the PML signal in G2 phase cells (Supplementary Fig. 2A and B). In interphase, the BLM signal also partly colocalized with the DNA repair markers RAD51, and 53BP1, likely in the previously observed repair foci (Supplementary Fig. 2C and D) (13,42). Strikingly, the G2 BLM foci disappeared shortly before the nuclear envelope breakdown (NEBD) (Fig. 2B) and the BLM signal became diffuse and dropped in intensity both with and without replication stress (Fig. 2C). In anaphase, the BLM-GFP protein was newly recruited to UFBs (Fig. 2B). Surprisingly, we also observed *de novo* formation of BLM foci on mitotic chromatin in late anaphase that persisted till the end of telophase, hereafter referred to as M-foci (Fig. 2B, see also Movie M2). To analyse the dynamic pattern of BLM-GFP localization, we tracked the mean intensity of individual foci over time, aligning all foci tracks at the NEBD. Averaging the foci intensity per condition revealed a slightly higher intensity of the G2 BLM foci in cells exposed to mild replication stress (APH) compared to the control (Fig. 2D). This detailed analysis further confirmed that BLM-GFP is released from chromatin upon NEBD, irrespective of replication stress, and is *de novo* recruited in anaphase.

**Figure 2.**
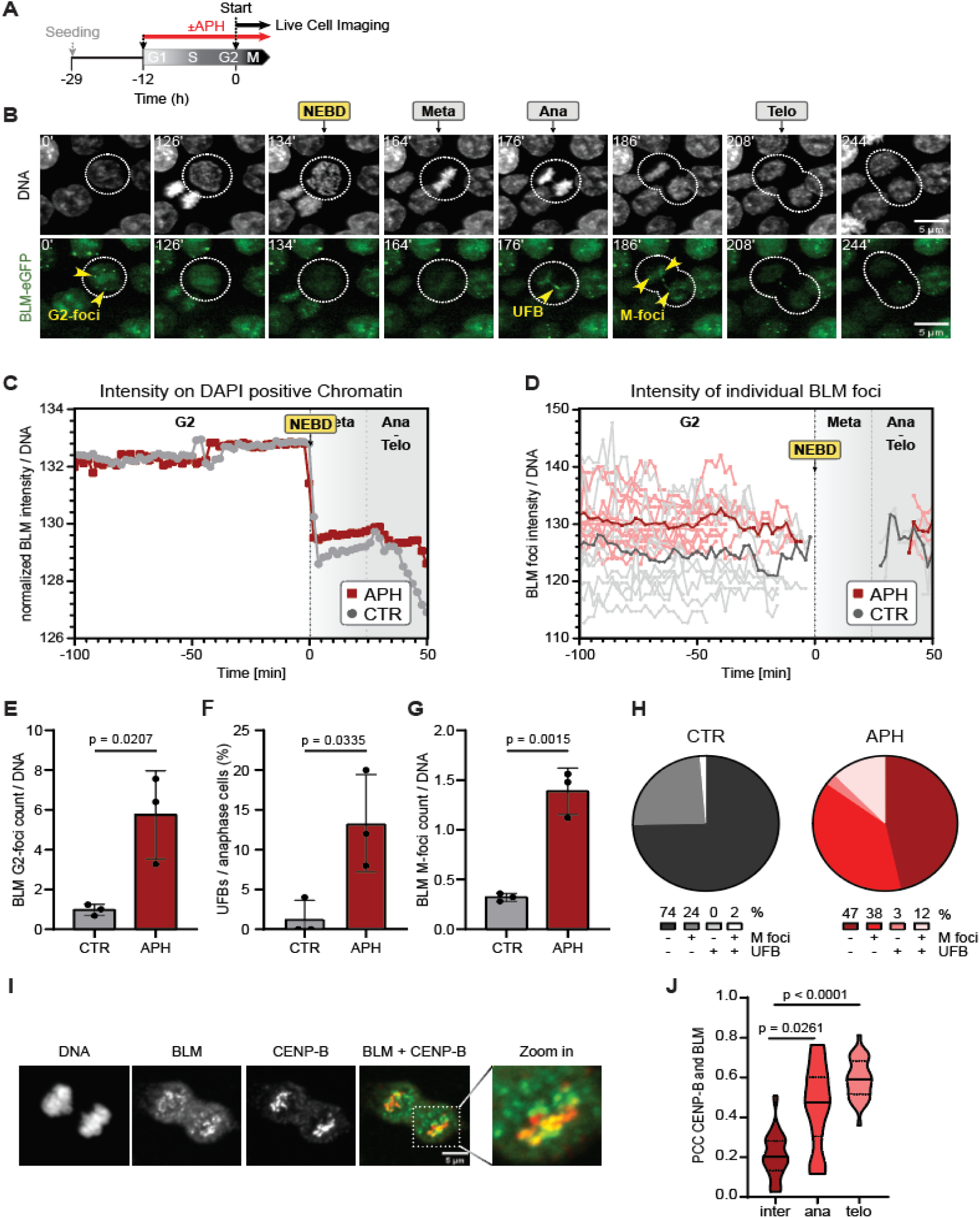
Endogenous BLM shows a bimodal recruitment to chromatin. **(A)** Experimental scheme. APH – aphidicolin. (**B**) Representative stills taken from live cell imaging of BLM^AID/GFP^ cells treated with 100 nM aphidicolin (APH), showing progression from G2 phase to mitotic exit with UFB and M-foci in ana-/telophase. White dotted circles indicate the cell of interest; yellow arrows mark BLM positive G2-foci, M-foci, or UFBs. NEBD: nuclear envelope breakdown; scale bar: 5 µm, the time (min) relative to the start of the movie is indicated on the top left corner. See Movie S1A. DNA was labeled with SPY 555. (**C**) Mean BLM-GFP intensity on chromatin over time tracked in individual cells (15 cells per biological replicate, n = 3) using TrackMate (Fiji plugin), aligned at NEBD (timepoint 0), and normalized to the untreated control (CTR). (**D**) Intensity of individual BLM-GFP foci over time, tracked as in (C) (5 cells per replicate, n = 3). Met: Metaphase, Ana: Anaphase, Telo: Telophase. (**E-G**) Percentage of cells with G2 foci (E), BLM positive UFBs (F), or M-foci (G). (**H**) Percentage of cells with no foci, only M-foci, only BLM positive UFBs, or both. Panels E-H show data from 25 cells per biological replicate (n = 3), mean ± SD is shown. For the analysis of BLM foci, a threshold was set based on APH-treated cells. The same threshold was used for G2- and M-foci. Live cell imaging was performed with a Lattice Light Sheet 7 microscope. (**I**) Representative microscopy images of BLM and CENPB co-localization in mitotic cells. (**J**) Pearson correlation coefficient of BLM and CENP-B signals. Mean and quantiles are shown as a black dotted line. 25 cells per biological replicate were evaluated, n = 3.

We next compared the number of BLM-foci and UFBs in control and APH-treated cells captured by live-cell imaging (Fig. 2E-G). Unstressed BLM^AID/GFP^ cells contained on average one G2 foci per cell, and after induction of mild replication stress, the number increased about 5-fold (Fig. 2E). Only 1 % of unstressed cells contained a visible UFB and less than one M-foci, while mild replication stress increased the fraction of UFB containing cells to 13 % on average, and the number of M-foci 5-fold (Fig. 2F and G). Additionally, mild replication stress increased the number of cells with co-occurring M-foci and UFBs 6-fold compared to untreated control (Fig. 2H). To further characterize the transient M-foci, we tested co-localization with PML or with the DNA repair markers γH2AX, RAD51 and 53BP1. In M-foci, BLM co-localized neither with PML nor with 53BP1 or RAD51, although there was a clear co-localization observed in G2 phase, suggesting that M-foci of BLM helicase represent a different physiological function (Supplementary Fig. 2A-D). Additionally, we observed almost no overlap between the BLM M-foci and γH2AX in non-treated mitotic cells and only a marginal increase of the Pearson correlation coefficient from 0.31 to 0.37 upon treatment with 100 nM APH (Supplementary Fig. 2E and F). Since the BLM helicase was recently proposed to contribute to centromere stability in mitosis (27,43), we stained the kinetochores with CENP-B antibody to evaluate the co-localization with the BLM signal (Fig. 2I and J). Strikingly, a large fraction of the BLM signal in the M-foci was next to, or overlapping with, the CENP-B signal in anaphase and telophase, while no overlap was observed in interphase, prophase, and metaphase (Fig. 2I and J, Supplementary Fig. 2G). Taken together, the dynamic localization patterns of the endogenous BLM reveal a *de novo* recruitment of the DNA helicase in mitosis to UFBs, as well as to novel M-foci.

### Both interphase and mitotic BLM functions contribute to the maintenance of genome stability

Given that BLM helicase is newly recruited to chromatin in mitosis, we asked whether restricting BLM depletion to M phase only would cause a similar phenotype as the continuous depletion across the entire cell cycle. Based on the cell cycle profile of cells synchronized by a single thymidine block, we selected the time point for cell phase-specific BLM depletion (Fig 3A and B, Supplementary 3A-D). Adding auxin 5 h after the release from thymidine block led to a gradual decrease of the BLM signal, with less than 10 % left at 10.5 h, when most cells reached mitosis (Fig. 3C and D). With this treatment, referred to as “M” depletion, cells undergo S phase in the presence of BLM helicase, while in mitosis BLM is depleted. Additionally, we treated the cells with auxin for the entire course of the experiment, hereafter referred to as “*S+M”* depletion (Fig. 3C and D). We also attempted to restrict the depletion of the BLM helicase to S phase only by washing out auxin 5 h after the release from the thymidine block. However, there was no detectable recovery of the BLM protein (Fig. 3C and D), and therefore, we did not continue with these experiments.

**Figure 3.**
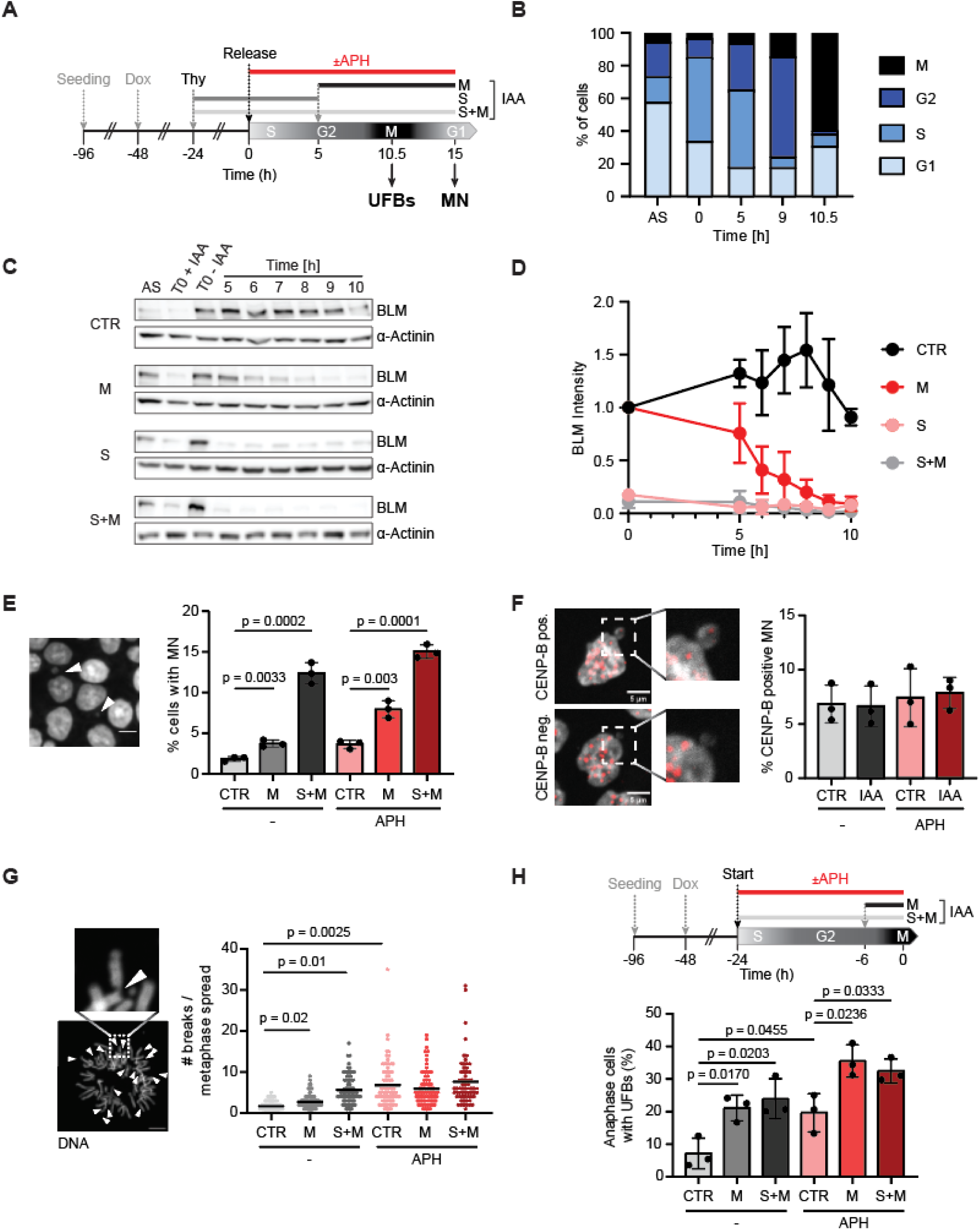
BLM depletion restricted to mitosis increases mitotic aberrations. **(A)** Experimental scheme. APH – aphidicolin, UFB - ultrafine bridges, MN – micronuclei, IAA – auxin, Thy – Thymidine, Dox – doxycycline. (**B**) Cell cycle profile of BLM^AID/GFP^ cells released from a single thymidine block. AS -asynchronous control. M phase was determined by staining with an MPM2 antibody, see Supplementary Fig. 3A-C. (**C**) Representative immunoblots of BLM in cells after release from single thymidine block with IAA added at specific timepoints as shown in panel A. α-actinin serves as loading control. CTR – control (no IAA), M – IAA added at 6 h after the release, S – IAA added at the synchronization start and washed out 6 h after the release, S+M – IAA added at the synchronization start, AS – asynchronous cells. (**D**) Quantification of the BLM abundance from panel C, normalized to the timepoint 0. (**E**) Percentage of cells in the subsequent G1 (15 h after release from the single thymidine block) with micronuclei (MN). Cells were treated as shown in (A), except that auxin was added 6 h after the thymidine release. APH – aphidicolin. Student’s t-test, mean ± S.E.M.; > 400 cells per replicate; 3 independent experiments. (**F**) Fraction of CENP-B positive micronuclei (MN) in asynchronous BLM^AID/GFP^ cells with or without treatment with IAA for 24 h. Student’s t-test, mean ± SD is shown, n = 3. (**G**) Chromosome spreads from cells harvested 10.5 h after the release from single thymidine block. Cells were treated as shown in (A), except that auxin was added 6 h after the thymidine release. Chromosome breaks were counted in 25 metaphase cells per replicate, 3 independent experiments, mean ± S.E.M. is shown. APH – aphidicolin. (**H**) Fraction of cells with PICH-positive UFBs in asynchronous cells treated for 24 h with aphidicolin. IAA was added together with aphidicolin (S+M depletion) or 6 h before harvesting (M depletion). Student’s t-test, mean ± SD is shown, three biological replicates.

Having established the mitotic depletion of BLM, we analysed its consequences on mitotic aberrations in cells released from thymidine block into media with or without APH. We first quantified micronuclei, which harbor broken or mis-segregated chromosomes (44). In the absence of replication stress, S+M depletion of BLM caused a 5-fold increase in micronucleated cells, while M depletion caused a two-fold increase. Replication stress led to an additional increase in micronucleation in all cases (Fig. 3E). 95 % of micronuclei contained a broken chromosome, as evidenced by the lack of staining with anti-CENP-B antibody, a marker of the inner kinetochore (Fig. 3F). Together, this suggests that the damage leading to micronuclei in these cells arises largely in S phase. Because the majority of the micronuclei contained a fragment of a broken chromosome, we next quantified DNA breaks and gaps on metaphase spreads (Fig. 3G). This further confirmed that primarily the S phase functions of BLM suppress the formation of chromosome breaks and gaps. In contrast, depletion of BLM only in mitosis led to a similar increase of anaphase cells with lagging chromatin or chromatin bridges, as depletion throughout the entire cell cycle (Supplementary Fig. 4A-B). Furthermore, the loss of BLM helicase in mitosis led to a two-fold increase in anaphase cells with PICH positive UFBs (Fig. 3H). Importantly, depletion over the entire cell cycle did not cause any further increase of UFBs in both asynchronous and synchronous cell populations (Fig. 3H, Supplementary Fig. 4C). Additionally, exposure of the cells to mild replication stress further elevated the formation of UFBs, but the effect of BLM depletion remained similar. Thus, we conclude that the BLM protein plays an essential role in M phase to maintain genomic stability independently of its function in S phase.

### Visualization of replication stress-induced UFBs in BLM-depleted cells reveals persistent sister chromatid linkages

To study the effect of BLM depletion on UFB resolution by live cell imaging, we additionally tagged the PICH protein with mCherry in the BLM^AID/GFP^ cells (Fig. 4A, Supplementary Fig. 5A-D). mCherry-positive cells were sorted by flow cytometry, and single cell clones were validated by Western blotting with antibodies against mCherry and PICH for homozygous tagging (Supplementary Fig. 6A). In the double-tagged cell line, the PICH-mCherry signal localized diffusely in the cytosol of interphase cells and rapidly re-localized to chromatin upon nuclear envelope breakdown, as previously reported (25,26) (Supplementary Fig. 6B and C, movie M3A-B). On mitotic chromatin, a weak PICH-mCherry signal was observed along all chromosomes. As soon as the cells entered anaphase, bright PICH-positive foci and UFBs appeared transiently in a subset of cells. Within these foci, and on UFBs, the PICH and BLM signals colocalized with high correlation (Fig. 4B and C, see also movie M3A-B). Notably, using antibodies for indirect immunofluorescence detection in the same cell, a strong PICH signal was observed along the BLM-positive UFBs, whereas the localization to M-foci was much less apparent, possibly due to high background signal observed with the PICH antibody (Fig. 4B, lower row). A similar depletion of the PICH signal on condensed chromosomes in immunofluorescence experiments has also previously been observed (e.g., (45,46)). We speculate that this might be either due to posttranslational modifications interfering with antibody binding or due to a general inaccessibility of the epitope within the condensed chromatin. Taken together, endogenous tagging of BLM and PICH revealed highly correlated mitotic signals both on UFBs and M-foci.

**Figure 4.**
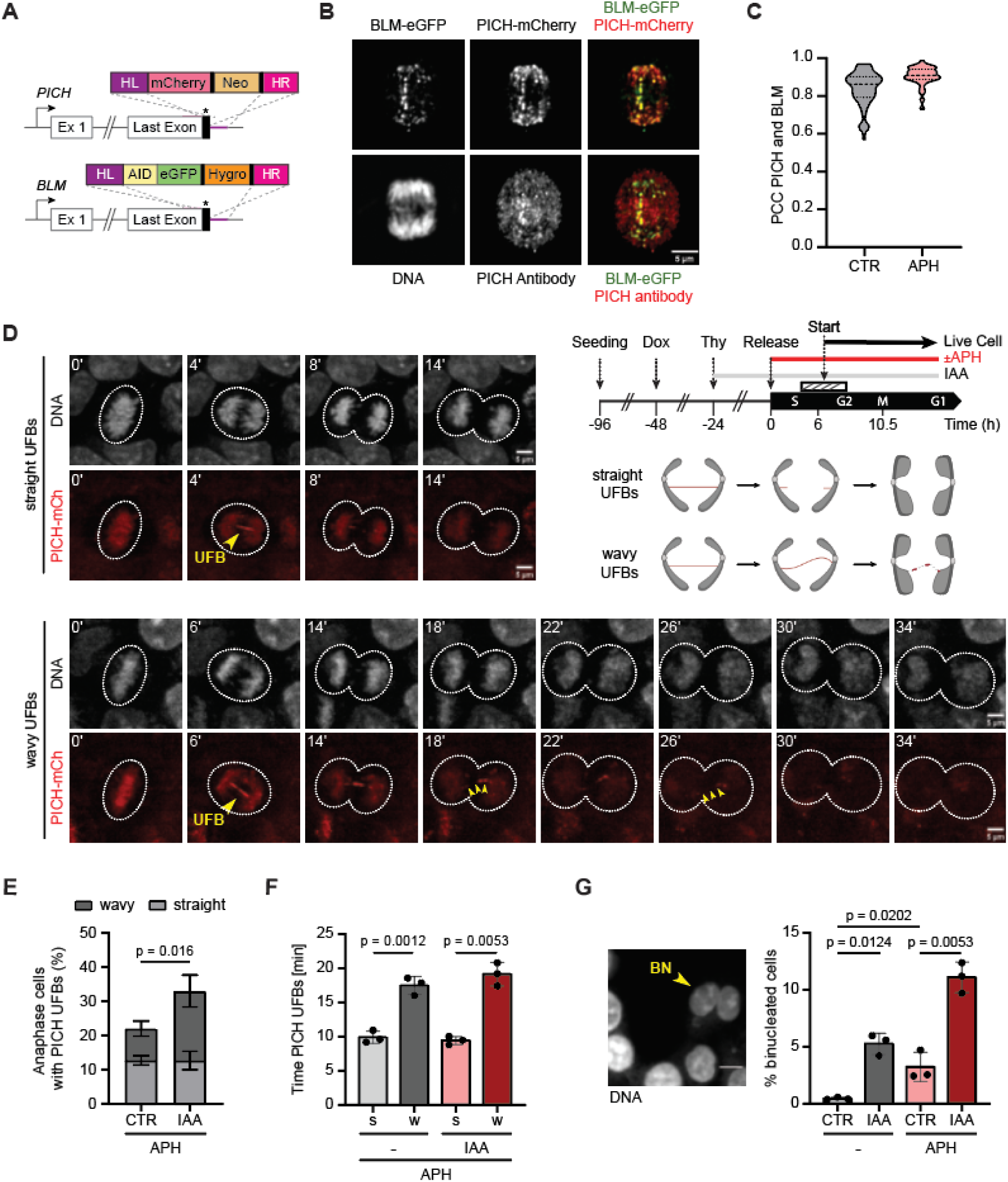
Visualization of replication stress-induced UFBs in BLM depleted cells reveals persistent sister chromatid linkages. **(A)** Schematic illustration of the gene editing strategy in BLM^AID/GFP^ cells. Imaging was started app. 6 h after the released from a single thymidine block (hashed bar). (**B**) Representative microscopy image of a fixed anaphase cell. BLM was visualized with the GFP tag, PICH was visualized either by an antibody or by direct mCherry fluorescence detection, DNA was stained with DAPI. (**C**) Pearson Correlation coefficient for the eGFP and mCherry signal in the nucleus. Data from 75 cells were pooled from 3 independent experiments for each condition. (**D**) Representative still images of cells synchronized and treated as in C and a schematic illustration of the live cell imaging strategy. The UFBs were visualized with PICH-mCherry, DNA was stained with SPY555. The cartoon depicts the two different fates of UFBs. Scale bar: 5 µm. APH – aphidicolin, UFB - ultrafine bridges, MN – micronuclei, IAA – auxin, Thy – Thymidine, Dox – doxycycline. (**E**) Quantification of the UFBs detected by live cell imaging (panel C). CTR – control, IAA – auxin, APH – aphidicolin. Student’s t-test, mean ± SD is shown, n = 3. (**F**) Duration of the PICH-mCherry signal along UFBs shown separately for “straight” (s) and “wavy” (w) UFBs. Student’s t-test, mean ± SD is shown, n = 3. For each replicate, 25 cells were tracked in live cell movies as shown in 4D. A detailed cell treatment scheme is provided in 4C. (**G**) Quantification of binucleated cells. Cells were fixed 15 h after release from a single thymidine block. IAA was added together with thymidine at the start of the synchronization and kept throughout the course of the experiment. Student’s t-test, mean ± S.E.M.; > 200 cells per replicate (n= 3). Scale bar, 10 µm.

Using the BLM^AID/GFP^/PICH^mCherry^ double-tagged cell line, we analysed the consequences of BLM depletion on UFBs by live cell imaging under mild replication stress. Consistent with the results obtained with fixed cells (compare Fig. 3H and Supplementary Fig. 4C), BLM depletion led to a small, but significant increase of mitotic cells with PICH positive UFBs (Supplementary Fig. 6). PICH localized to UFBs both in the presence and absence of BLM, further supporting its upstream role in the recruitment of the BLM helicase and other proteins to UFBs, as previously suggested (20,46). Interestingly, BLM depletion also prolonged the lifetime of UFBs by about 4 minutes on average (Supplementary Fig. 6E), whereas the time for the passage from the start of anaphase to the mitotic exit remained unchanged (Supplemental Figure 6F). Careful tracking of anaphase across multiple time frames revealed two distinct classes of PICH-positive UFBs (Fig. 4D). In the first class, UFBs appear as a straight line of the PICH signal, which disappears from the middle to the sides within a few minutes (Fig. 4D, upper panel, “straight” UFBs, Movie M4A-B). In the second class, the originally “straight” UFBs became “wavy” in the later stages of mitosis, and the PICH signal dissipated into a small number of foci along the path of the UFB before disappearing (Fig. 4D, lower panel, Movie M5A-B).

Intriguingly, upon depletion of the BLM helicase, only the fraction of the “wavy” UFBs increased more than two-fold compared to BLM positive cells, while the “straight” UFBs appeared unaffected (Fig. 4E). Strikingly, the PICH signal on the “wavy” UFBs was present twice as long than on the “straight” UFBs (20 vs. 10 min, Fig. 4F). Thus, most of the “straight” UFBs were dissolved before chromatin de-condensation started, while “wavy” UFBs were still observed in telophase (compare Fig. 4D, upper and lower panel). It is important to consider that the loss of the PICH signal towards the end of anaphase does not necessarily mean that the linkage between the sister chromatids has been resolved. We therefore hypothesized that these unresolved DNA bridges might interfere with cytokinesis, giving rise to binucleated cells. Indeed, we observed a significant increase in binucleated cells upon BLM depletion in fixed cells, which was further exacerbated by mild replication stress (Fig. 4G). These results suggest that the loss of BLM function leads to increased occurrence of “wavy” ultra-fine bridge structures, which take longer to resolve. Unresolved “wavy” UFBs may thus contribute to the formation of binucleated cells, leading to whole genome doubling and potentially promoting aneuploidy and cancer.

### Single-cell sequencing uncovers genomic instability and whole genome doubling after BLM depletion

Next, we performed single-cell whole genome sequencing (scWGS) to define genomic copy number alterations (CNAs) in BLM-depleted cells that progressed through one cell cycle in the presence of mild replication stress. To this end, we synchronized the cells by a single thymidine block and released them into media with 100 nM APH (Fig. 5A). The BLM helicase was either depleted throughout the entire experiment (S+M), or only during M phase. At the release, we added EdU to label DNA in replicating cells, and after 6 h, the CDK4/6 inhibitor palbociclib to trap the cells in the subsequent G1 to obtain homogenous populations for scWGS. Flow cytometry confirmed that more than 85 % of the cells were trapped in G1, and 76 to 87 % of these cells underwent replication during the 24 h (Fig. 5B).

**Figure 5.**
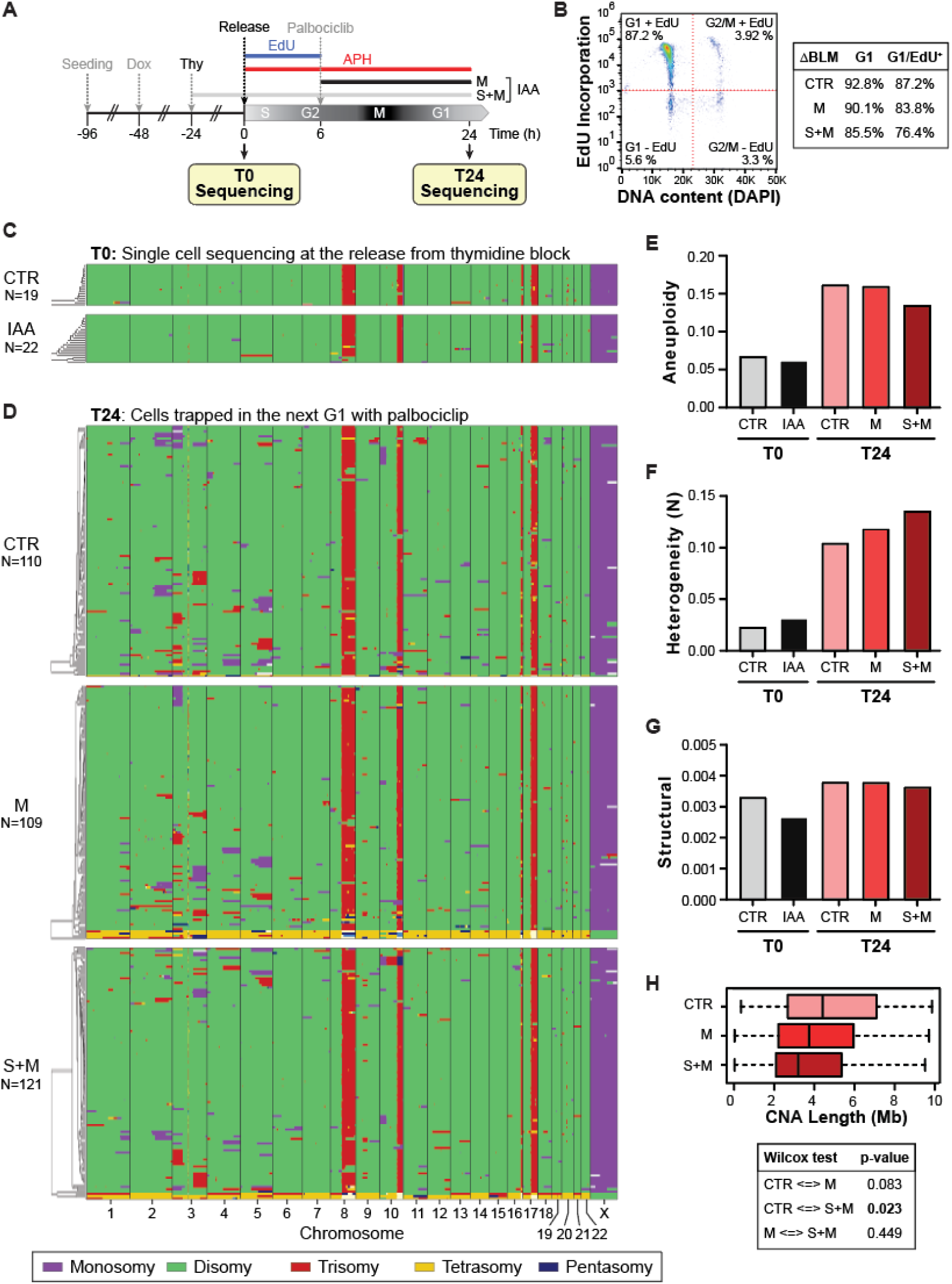
Single-cell sequencing reveals genomic instability after replication stress. **(A)** Schematic illustration of the experimental strategy. APH – aphidicolin, IAA – auxin, Thy – Thymidine, Dox – doxycycline. (**B**) Flow cytometry analysis of EdU incorporation in cells treated as shown in (A). The table on the right specifies the fraction G1 cells scoring positive for EdU. (**C**)(**D**) Genome-wide copy number plots generated by the Aneufinder algorithm. Cells treated as shown in (A) were subjected to low coverage scWGS. Sequence reads were aligned to the human reference genome (GRCh38/hg38) and averaged over 1 MB non-overlapping bins. Note that the structural chromosomal changes within t(8p;18q), 10q+, t(9q;16p-) 16q and 17q, as well as chromosome Y loss are characteristic of HCT116. (**E**-**G**) Aneufinder analysis of the level of aneuploidy, heterogeneity and structural aberrations (see Methods for details). (**H**) Distribution of CNA fragment length. Results from a Wilcoxon test are shown below the graph.

As expected, genomes of cells analysed at the time of release (T0) showed only a low level of chromosomal aberrations, confirming that HCT116 is a chromosomally stable cell line (Fig. 5C) (47). In contrast, passage through a one cell cycle only in the presence of mild replication stress caused a significant increase of genomic aberrations defined by the number of CNAs (Fig. 5D). Quantification of aneuploidies and CNAs using the Aneufinder algorithm (36) revealed a two-to three-fold increase of aneuploidy and sample heterogeneity, whereas structural aberrations remained largely unchanged (Fig. 5E-G). Although BLM depletion did not significantly alter aneuploidy or the frequency of structural aberrations (Fig. 5E and G), it modestly increased sample heterogeneity (Fig. 5F) and led to a pronounced accumulation of tetraploid cells (Fig. 5D marked in yellow). To robustly quantify the heterogeneity of CNAs, we removed all technical artifacts (see Methods) and normalized the read counts of T24 samples to the median of T0 controls, which were harvested prior to the release into S phase (Supplementary Fig. 7A-C). To exclude CNAs involving entire chromosome arms, we filtered the dataset to retain only CNAs shorter than 10 Mb (see “Material and Methods” section). Although this refined analysis revealed no statistically significant differences in the heterogeneity scores among T24 samples (Supplementary Fig. 7D-F), BLM depleted samples displayed a significant reduction of the median CNA fragment length from 4.4 Mb to 3.2 Mb (Fig. 5H). Collectively, the scWGS data indicate that BLM depletion leads to an accumulation of tetraploid cells and a shift towards shorter-sized CNA fragments, supporting the notion that, in the absence of BLM, a subset of replication intermediates is resolved through an alternative pathway or not resolved at all.

## Discussion

The BLM helicase supports many different facets of DNA metabolism. While the BLM functions during the S and G2 phase in DNA replication and homologous recombination-mediated repair have been extensively studied and well established, its roles in mitosis remain poorly understood. By establishing a cell line that allows live cell imaging of the BLM and PICH helicases along with an acute and efficient BLM depletion, we demonstrate that BLM has distinct functions during M phase that are essential for maintaining genome stability. Live cell imaging revealed a remarkably dynamic localization of the BLM and PICH helicases throughout the cell cycle. Most strikingly, we observed that the BLM protein was completely released from chromatin shortly before nuclear envelope breakdown and regained access only later in anaphase, where it accumulated at ultrafine bridges as well as in a few discrete mitotic foci (Fig. 2B-D). The release of BLM coincided precisely with the high activity of the protein kinase CDK1, which is required to trigger the entry into mitosis, chromosome condensation and nuclear envelope breakdown. Intriguingly, Western blotting and proteomics experiments showed that BLM becomes heavily modified by CDK1-, PLK1-, and MPS1-dependent phosphorylation as cells enter mitosis (48). In line with our observation of dynamic BLM behaviour during mitosis, the phosphorylation is posed to change its subcellular localization, as the hyperphosphorylated BLM was exclusively contained in the soluble nucleoplasm, while only the unmodified protein remained in the insoluble nuclear matrix (3). This suggests that the phosphorylation initiates the release of BLM from chromatin, PML bodies, and the DNA repair foci shortly before the nuclear membrane is disassembled. Additionally, PLK1-dependent phosphorylation also prevents BLM from binding to centromeres, thereby avoiding premature unwinding of the centromeric DNA during the early stages of mitosis, an activity the authors termed the “centromere protection pathway” (49). How the BLM helicase regains its DNA binding activity later during anaphase to localize to UFBs and the mitotic foci remains poorly understood. While CDK1 activity sharply declines at the onset of anaphase (50), the phosphorylation status of mitotic proteins is additionally regulated by protein phosphatases. For example, protein phosphatase 1 (PP1) together with RIF1 becomes recruited to PICH-containing UFBs to regulate the assembly of the UFB resolution complex (51). Given the extensive modification of UFB binding proteins by multiple mitotic kinases and SUMO E3 ligases, our newly established cell line may serve as a powerful tool to further dissect this highly dynamic regulatory network.

Another intriguing observation from our live-cell imaging is the accumulation of the BLM helicase during ana- and telophase in a few mitotic foci that also stain positive for the PICH helicase (Fig. 2B, Fig. 4B). By examining their co-localization with several prototypic marker proteins, we demonstrate that the largest subset is found at centromeres. Earlier reports also demonstrated a focal accumulation of PICH near centromeres throughout mitosis (25,52,53). However, in contrast to PICH, BLM localization at the centromeres is likely more transient, starting only after anaphase onset, which is critical, as uncontrolled BLM activity in metaphase cells appears to compromise centromere stability (49,54). While some of these foci may simply reflect remnants of resolved UFBs, they may also point to a new function of BLM and PICH at a subset of centromeres. Human centromeres are regions that replicate later in the S phase and are composed of long, AT-rich repetitive sequences with a tendency to form non-B-DNA structures and DNA:RNA hybrids (55). Their structure makes centromeres inherently prone to DNA breakage, which is exacerbated under replication stress (56). Recent proteomic studies revealed an enrichment of BLM and other components of UFB resolving protein complexes on centromeric DNA, potentially acting to counterbalance its fragility (43). It therefore will be interesting to elucidate, whether and how the BLM helicase accumulating in select CENP-B positive foci late in mitosis contributes to the suppression of DNA breaks in pericentric DNA.

Using rapid, auxin-induced depletion we show that passage through a single cell cycle in the absence of the BLM helicase induces genomic instability, as evidenced by a steep increase of micronucleated cells, chromosomal breaks, and anaphase cells with UFBs (Fig. 3, Supplementary Fig. 4). Of note, such acute depletion eliminates selection of suppressor mutations commonly observed in BLM deficient cells (57), and which may severely confound the effects of BLM depletion on genome stability. The rapid depletion further allowed us to restrict BLM loss to a single mitosis, enabling us to study individually the contribution of early and late BLM functions to the maintenance of genome stability. Strikingly, mitotic depletion of BLM increased the number of UFBs to similar levels as in cells with continuous depletion over the entire cell cycle (Fig. 3H, Supplementary Fig. 4C). This suggests that the early BLM functions in S and G2 phase contribute only little to the suppression of UFB formation and that the accumulation of UFBs was mainly due to an impaired resolution of the sister chromatid linkages. Surprisingly, also lagging chromosomes increased to similar levels in both depletion schemes, whereas the level of micronucleation was higher in cells with continuous BLM depletion (Fig. 3E, Supplementary Fig. 4A and B). Together with the observation that most micronuclei stained negative for the CENP-B protein, we conclude that micronucleation is driven mostly by the early BLM functions in the S and G2 phase of the cell cycle that likely suppress the breakage of chromosomes and the formation of acentric chromosome fragments. In this context, it is interesting to note that micronuclei may form not only upon exit from mitosis, when the nuclear envelope reforms, but also during S phase (58). Consistent with our interpretation, fibroblasts derived from patients with Bloom syndrome were found to expel micronuclei at a high rate during S phase (12).

Strikingly, live-cell imaging revealed the existence of two different classes of UFBs, which can be distinguished according to their morphology and dynamics as short-lived, “straight” and long-lived “wavy” UFBs. While the proportion of cells exhibiting “wavy” UFBs clearly increased upon BLM depletion, the frequency of “straight” UFBs remained unchanged. The observation that all “straight” UFBs are rapidly resolved irrespective of BLM depletion (Fig. 4F) suggests the involvement of a redundant pathway in their resolution. The persistence of the “wavy” UFBs moreover suggests that their underlying structure differs from those of straight UFBs, making them resistant against cleavage or resolution by the back-up pathways. Additional experiments will be required to determine, how “straight” UFBs differ structurally from “wavy” UFBs, whether they are induced by distinct perturbations, and if they arise at different genomic loci.

To further investigate the consequences of acute BLM depletion on genome stability, we subjected the cells to scWGS. Similar to an earlier report (59), passage through a single cell cycle in the presence of low doses of APH caused the accumulation of a significant number of CNAs compared to synchronized cells harvested just before the release into S phase. However, except for the induction of a low number of tetraploid cells, and reduced CNA size, BLM depletion did not substantially increase the overall frequency of CNAs in our model system (Fig. 5). These results were surprising, given the synergistic effects of replication stress and BLM depletion on micronucleation, UFB frequency, as well as binucleation. The pronounced increase in DNA breaks following BLM depletion (Fig. 3G) along with the well-documented cancer predisposition associated with Bloom syndrome, raises the question of why acute BLM loss did not lead to detectable genomic instability in our scWGS analysis. One possible explanation is that additional cell divisions may be necessary for accumulating measurable genome rearrangements to become established or that the most prevalent genomic alterations remain undetectable due to the limitations of our short-read, low-coverage genome sequencing. Moreover, we hypothesize that mild replication stress alone induces such a high load of CNAs that the potentially more subtle contributions of BLM depletion remain obscured. Recently, high coverage whole genome sequencing revealed a strong increase of interchromosomal translocations in p53-deficient cells where PICH was depleted for three consecutive days (52). Similarly, low coverage scWGS revealed a modest increase of CNAs in knock-in cells harbouring patient-derived BLM mutations (60). Taken together, we conclude that acute loss of BLM during one cell cycle does not drive the immediate accumulation of CNAs in cells challenged with mild replication stress. Instead, additional rounds of DNA damage processing over multiple cell cycles may be required to repair the lesion, leaving behind scars in the DNA. Complementary experiments will be required to fully evaluate the genomic consequences of acute BLM depletion and to understand the molecular mechanism that underlie the altered CNA size distribution. Collectively, the detection of tetraploid cells in our scWGS analysis (Fig. 5D) together with the accumulation of binucleated cells following acute BLM depletion during a single cell cycle (Fig. 4G) provide evidence that the absence of BLM depletion leads to long-lived inter-sister chromatids linkages that impair cytokinesis. These results are consistent with the accumulation of binucleated cells in chronically BLM-deficient patient cells (20) or upon siRNA-mediated depletion of BLM, PICH or RIF1 (26,61). In our live-cell imaging, we observed a subset of “wavy” UFBs that persist long into telophase, coinciding with chromatin decondensation (Fig. 4D). Although we could not further track these bridges due to the release of the fluorescent marker proteins from the UFBs, they likely triggered the formation of binucleated cells, as DNA trapped at cleavage furrow interferes with abscission (62). In support of this idea, the post-mitotic cells with “wavy” UFBs often remained tightly connected to each other and showed symmetric sister-nuclei, a typical feature of binucleated cells. Our observations indicate that, in addition to unresolved chromatin bridges (62), persistent ultrafine bridges composed of chromatin-free DNA linkages between sister chromatids can impair abscission, leading to daughter cells that remain connected by stable intercellular canals. These findings underscore the essential role of BLM during mitosis in ensuring accurate segregation of chromatin into daughter cells and suggest that the mitotic BLM function directly supports genome stability by actively suppressing whole genome doubling, a recognized early and frequent driver of tumorigenesis.

## Data availability

scWGS data has been deposited at the European Nucleotide Archive (ENA) and can be accessed using the following ID: **PRJEB88724**

## Supplementary data

**Supplementary Figure 1.**
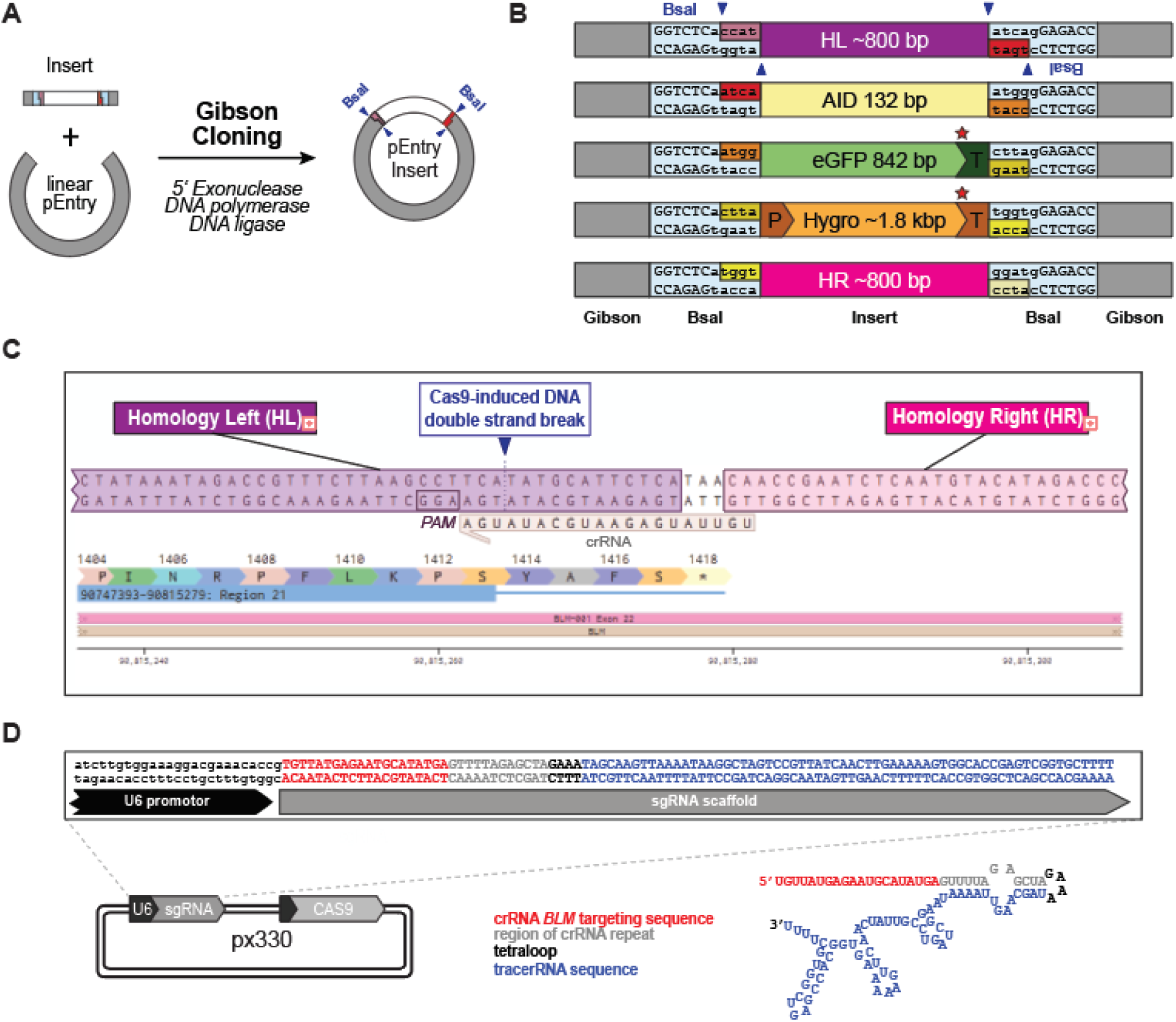
CRISPR/Cas9 gene editing of BLM for endogenous AID-eGFP tagging. **(A)** Schematic illustration of the entry plasmid cloning. (**B**) Individual inserts of the pEntry vectors. HL – homology left, HR – homology right, P – promotor, T – terminator, Gibson overhangs used for the Gibson cloning are shown in gray. Red stars indicate stop codons, blue arrows indicate cuttings sites of the BsaI restriction endonuclease. See Material and Methods for details. (**C**) Benchling view of the genomic *BLM* locus around the stop codon depicting the design of the sgRNA. (**D**) Schematic of the px330 plasmid with the cloned sgRNA targeting *BLM* and providing Cas9.

**Supplementary Figure 2.**
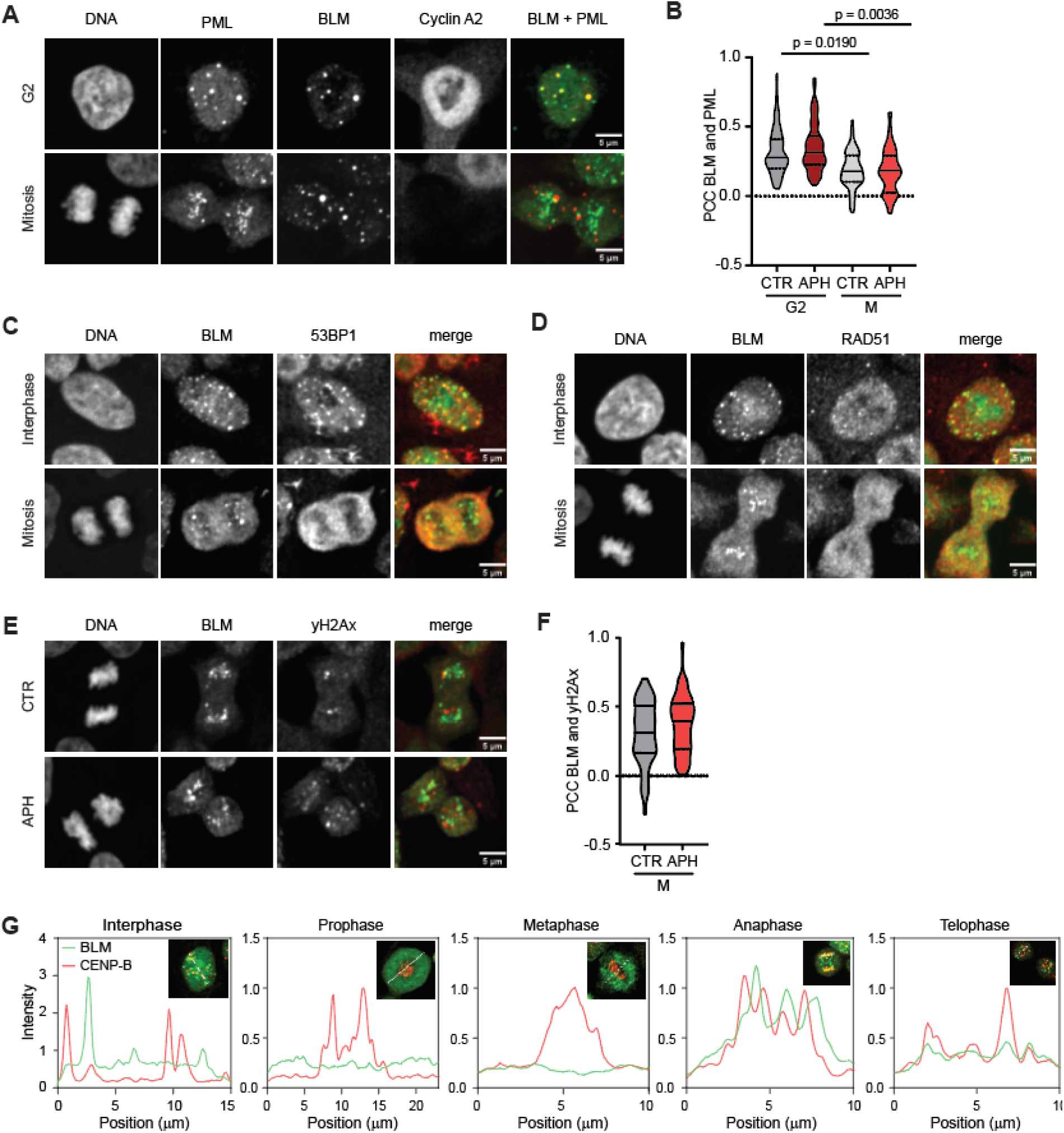
Characterization of G2 and M specific BLM foci. **(A)** Representative microscopy images of BLM and PML co-localization in G2 and M cells. (**B**) Pearson correlation coefficient of BLM and PML signals in G2 and M cells. Mean and quantiles are shown as a black dotted line. 25 cell per biological replicate were evaluated, n = 3. (**C**) Representative microscopy images of BLM and 53BP1 co-localization in interphase and M cells. (**D**) Representative microscopy images of BLM and RAD51 co-localization in interphase and M cells. (**E**) Representative microscopy images of BLM and γH2AX co-localization in anaphase cells. (**F**) Pearson correlation coefficient of BLM and γH2AX signals in anaphase cells. Mean and quantiles are shown as a black dotted line. 25 cells per biological replicate were evaluated, n = 3. (**G**) Co-localization of BLM and CENPB in different cell-cycle phases. The fluorescence intensity is plotted along an arbitrary line randomly placed across the cell nucleus (see white dotted line in insets).

**Supplementary Figure 3.**
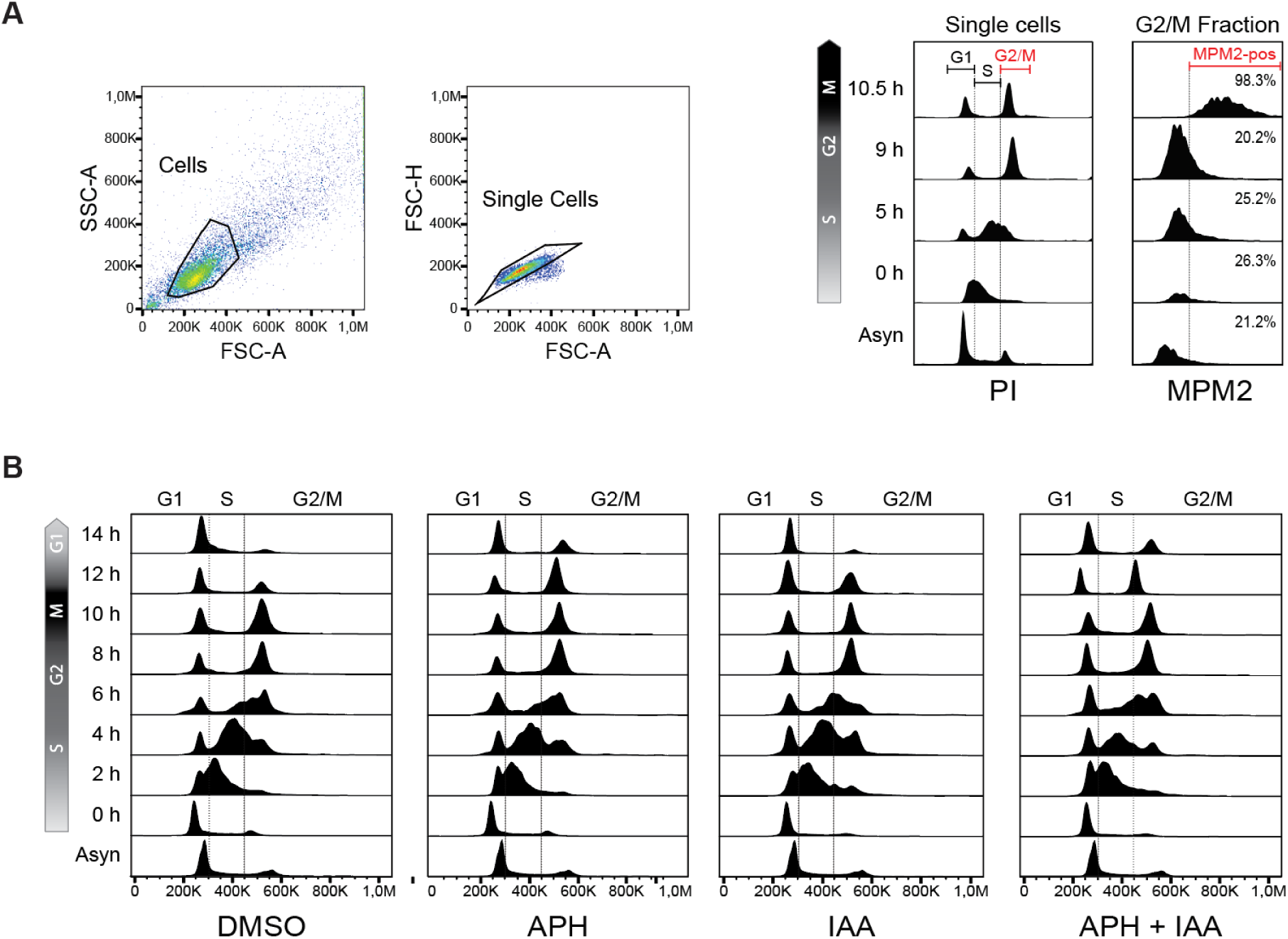
Cell cycle phase specific depletion of BLM helicase. **(A)** Flow cytometry analysis for determining cell cycle profiles of the BLM^AID/GFP^ cells released from single thymidine block (see Fig. 3A). G2/M cells were gated as shown and analyzed for binding the MPM2 antibody, which specifically recognizes mitotically phosphorylated proteins. (**B**) Cell cycle analysis of BLM^AID/GFP^ cells released from a single thymidine block into media containing DMSO, APH, IAA or a combination of APH and IAA. Note that IAA was added together with thymidine at the start of the synchronization (“S+M” depletion). IAA – auxin.

**Supplementary Figure 4.**
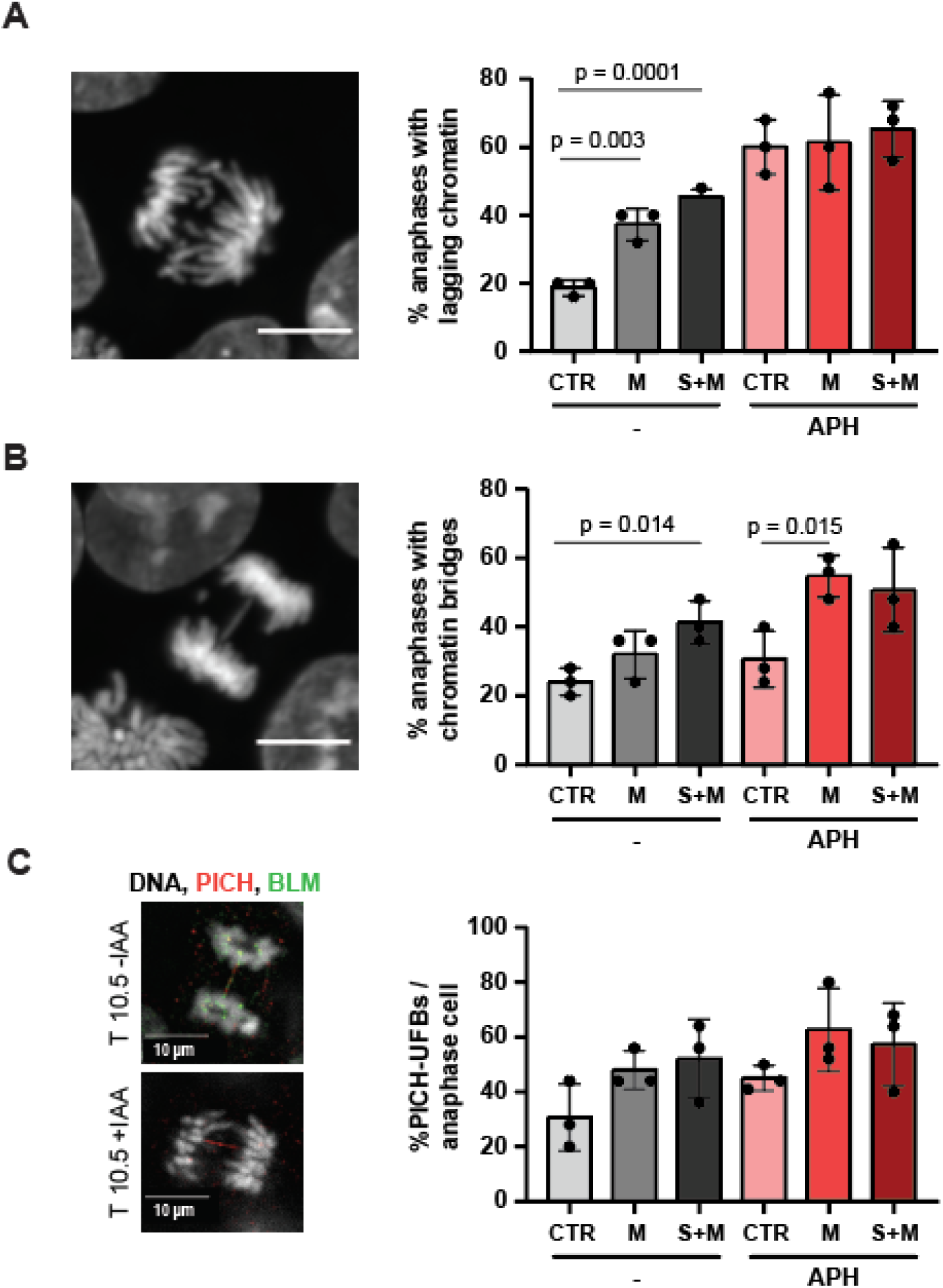
Cell cycle-specific depletion of BLM increases the frequency of mitotic aberrations. (**A-C**) Quantification of cells with lagging chromatin, chromatin bridges or PICH positive UFBs in control and BLM depleted cells. Cells were synchronized with a single thymidine block and released into media with or without 100 nM aphidicolin, as shown in Fig. 3A except that IAA was added 6 h after the thymidine release. 25 anaphase cells were counted in each of three independent experiments. Student’s t-test, mean ± SD is shown.

**Supplementary Figure 5.**
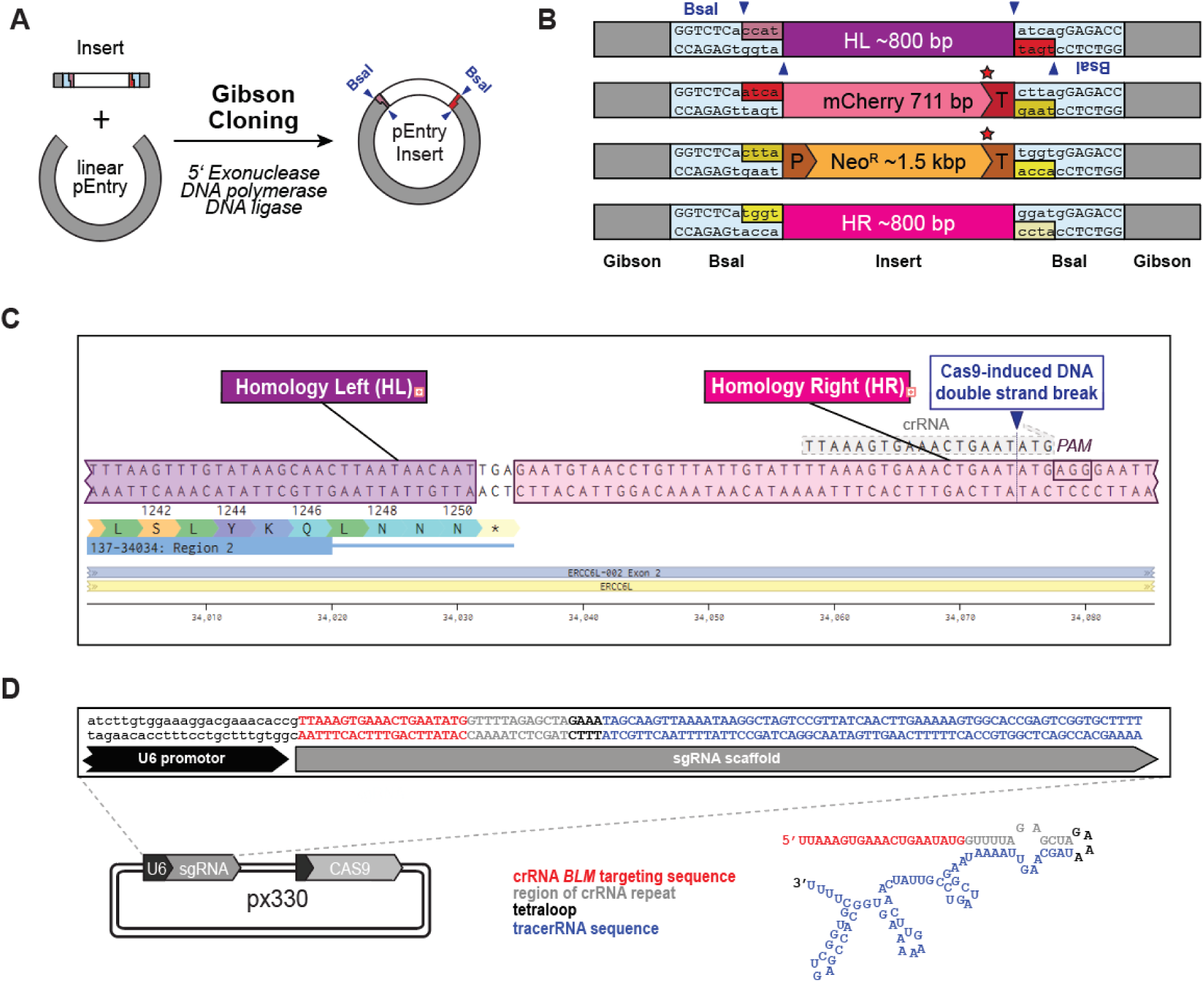
CRISPR/CAS9 gene editing of endogenous PICH for mCherry tagging. **(A)** Schematic illustration of the entry plasmid cloning. (**B**) Inserts of the individual pEntry vectors used for the assembly of the homology donor for PICH tagging (see Fig. 1A and Materials and Methods for details). HL – homology left, HR – homology right, P – promotor, T – terminator, Gibson overhangs used for the Gibson cloning are shown in gray. Red stars indicate stop codons, blue arrows indicate cuttings sites of the BsaI restriction endonuclease. See Material and Methods for details. (**C**) Benchling view of the genomic *PICH* locus around the stop codon depicting the design of the sgRNA. Note that in the right homology arm the protospacer adjacent motif (PAM) was mutated to AGA to prevent cutting by Cas9. (**D**) Schematic of the px330 plasmid with the cloned sgRNA targeting *BLM* and providing Cas9. The resulting px330 was transfected together with the linearized homology donor into the BLM^GFP/AID^ HCT116 cells.

**Supplementary Figure 6.**
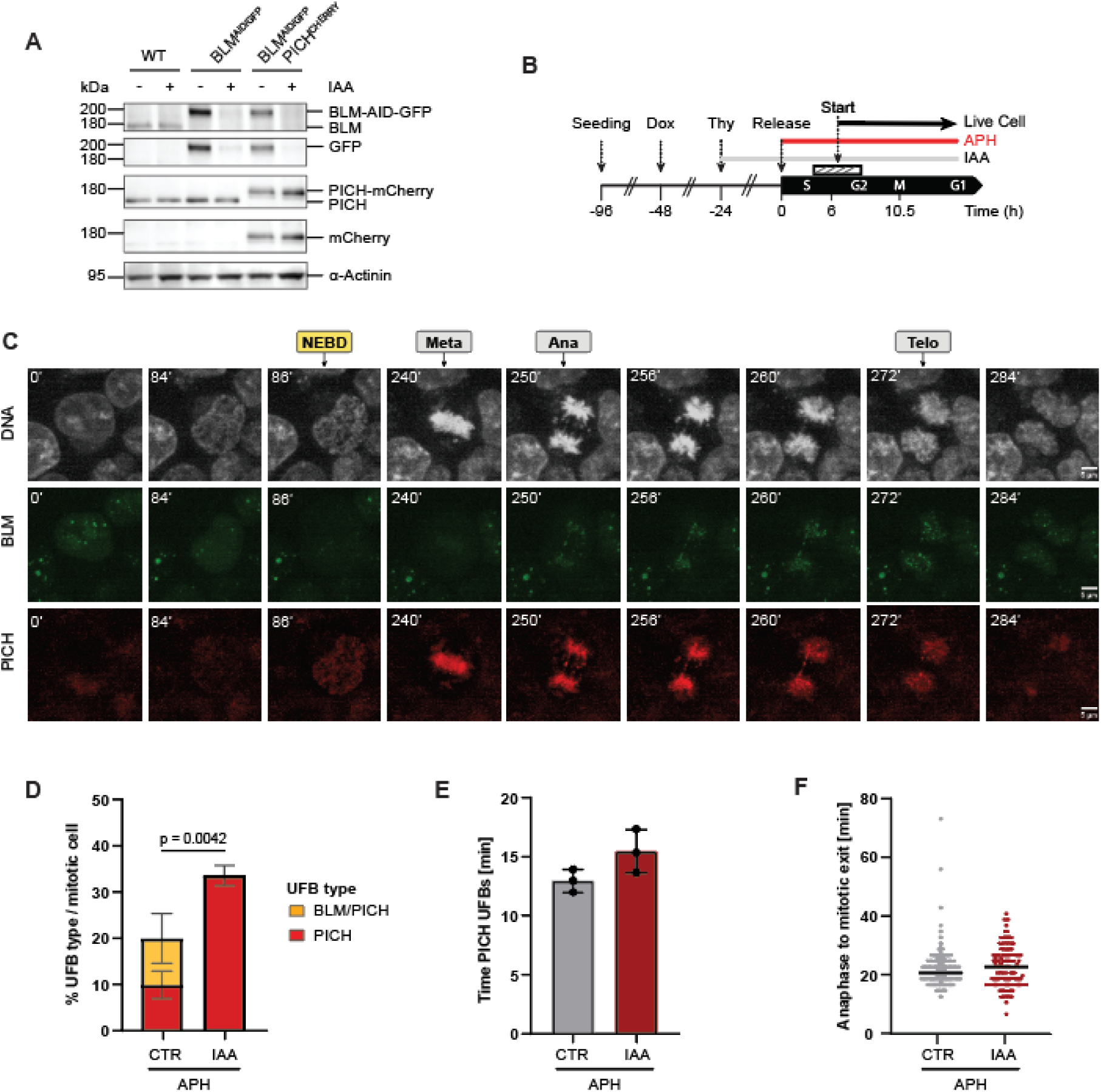
Validation of mCherry tagging of the endogenous PICH protein in BLM^AID/GFP^ cells. **(A)** Detection of untagged and tagged versions of BLM and PICH in the parental HCT116 cells, BLM^AID/GFP^ cells or BLM^AID/GFP^ with mCherry tagged PICH. Membranes were blotted with antibodies raised against BLM, GFP, PICH, mCherry or α-Actinin, respectively. α-Actinin served as a loading control. (**B**) Experimental scheme for the live cell microscopy experiment shown in (C). Imaging was started app. 6 h after the released from a single thymidine block (hashed bar). APH – aphidicolin, UFB - ultrafine bridges, MN – micronuclei, IAA – auxin, Thy – Thymidine, Dox – doxycycline. (**C**) Representative microscopy images from movie M3A-B. Movies were acquired from HCT116 cells expressing AID/GFP tagged BLM and mCherry tagged PICH treated as shown in (B). Labels indicate cells at nuclear envelope breakdown (NEBD), in metaphase (Meta), at the start of anaphase (Ana) or at the end of telophase (Telo). DNA was labeled with SPY 555, BLM and PICH were visualized via the fluorescence of eGFP and mCherry, respectively. (**D**) Fraction of anaphase cells with UFBs. More than 30 cells per biological replicate were analysed. Student’s t-test, mean ± SD is shown, n = 3. IAA – auxin, APH - Aphidicolin. The Student’s t-test was calculated for the comparison between all PICH positive UFBs in control and IAA treated cells including also PICH/BLM double positive UFBS. (**E**) Average time for which the PICH UFBs persist. At least 20 anaphases in each biological replicate (n = 3) were analysed. Student’s t-test, mean ± SD is shown. (**F**) Time cells spend from beginning of anaphase until mitotic exit. Mitotic exit was defined here as the time, when DNA appeared to be fully decondensed, as indicated in (A). More than 200 cells per replicate were evaluated. Student’s t-test, mean ± SD is shown, n = 2.

**Supplementary Figure 7.**
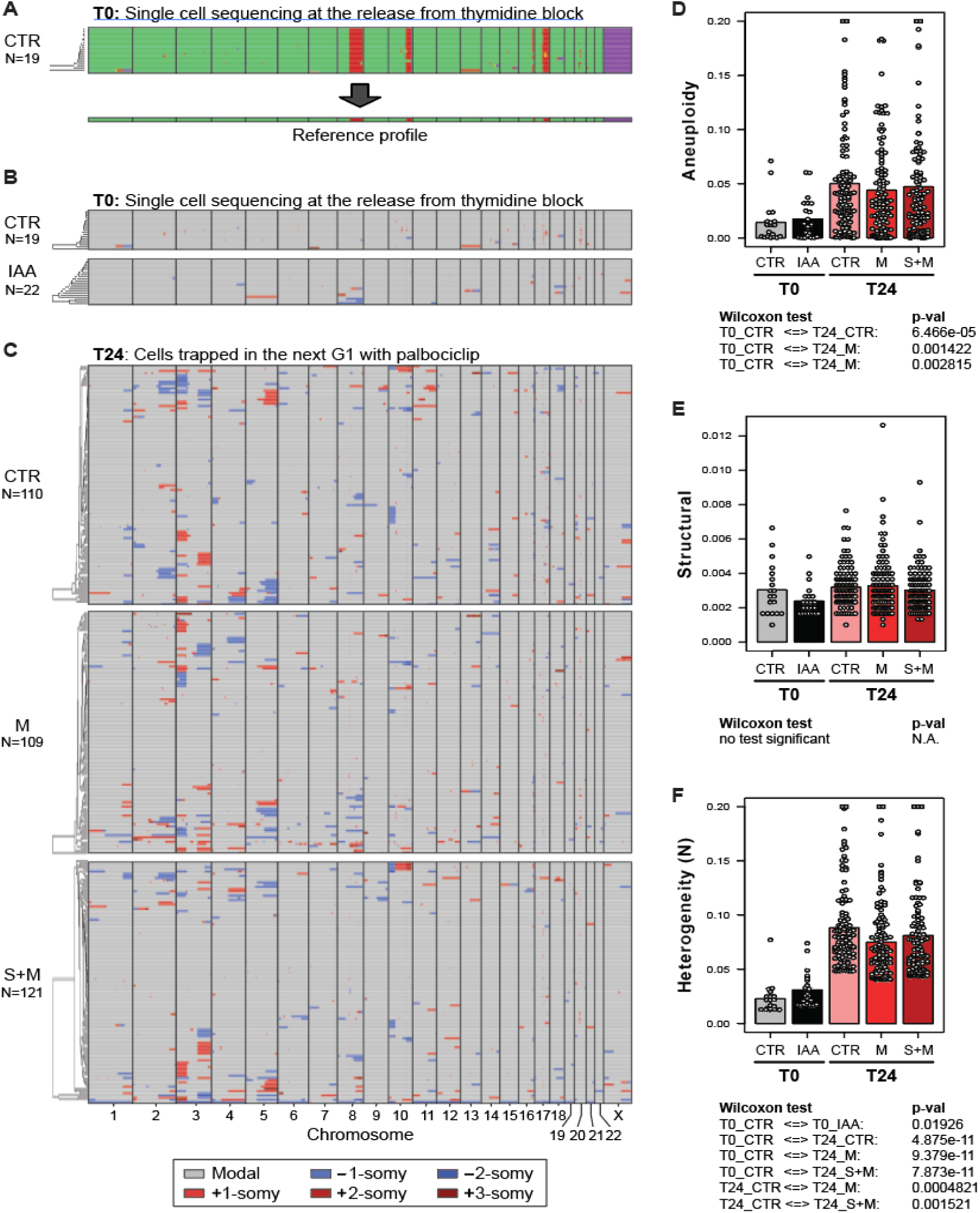
Re-analysis of scWGS data based on the alignment to T0 controls. **(A)** Schematic illustration how the T0 controls were median averaged to obtain the reference profiles for DMSO and IAA treated samples, respectively. (**B**) (**C**). Genome-wide copy number plots generated by the Aneufinder algorithm. Data shown in Fig. 5C and D was re-analysed. Sequence reads were aligned to the median average profiles of *“T0 – IAA”* and “T0 + IAA” and averaged over 1 MB non-overlapping bins. (**D**-**F**) Aneuploidy, heterogeneity and structural aberrations scores after re-analysis of the scWGS data (see Methods for details).

## Acknowledgements

We thank Masato Kanemaki for providing HCT116 cells expressing TIR1, Michael Taschner and Stephan Gruber for providing advice and reagents for the modular cloning strategy and Andrea Tijhuis for help with cell sorting and library preparation. Biorender was used to prepare the graphical abstract and Figure 4D.

## Funding

This project was funded by the FOR2800 Chromosome Instability: „Cross-talk of DNA replication stress and mitotic dysfunction” (DFG FOR 2800) to ZS, MR, by the European Research Council (ERC) Synergy Grant (GA 855158) to IMT, and a Dutch Cancer Society Grant (2017-RUG-11457) to FF.

## Author contribution

Conceptualization, M.R. and Z.S.; Methodology, M.R., Z.S.; Investigation, T.H., A.W., A.T., F.M., K.V., M.L., I.I.G., R.W. and F.F.; Writing – Original Draft M.R. and Z.S.; Writing – Review & Editing, all authors; Funding Acquisition, M.R., Z.S. and I.T.; Supervision, M.R., Z.S., I.T., B.W. and F.F.

## Conflict of interest statement

FF is Chief Scientific Director of iPsomics, a startup business that provides scWGS services to the academic community. All other authors declare no conflict of interest.

